# Comprehensive Analysis of CRISPR Base Editing Outcomes for Multimeric Protein

**DOI:** 10.1101/2022.06.20.496792

**Authors:** Meha Kabra, Mariya Moosajee, Gregory A. Newby, Kaivalya Molugu, Krishanu Saha, David R. Liu, Bikash R. Pattnaik

## Abstract

Point mutations in the *KCNJ13* gene cause an autosomal recessive, childhood blindness, Leber congenital amaurosis (LCA16) due to a loss-of-function Kir7.1 channel. In the present study, we investigated the etiology of LCA16 caused by a *KCNJ13* missense mutation (c.431T>C, p.Leu144Pro) and explored the activity of two cytosine base editors mRNAs (CBEs, BE4max-WTCas9, and evoCDA-SpCas9-NG) as a proof-of-concept therapeutic option. We observed the *KCNJ13*-related retinopathy phenotype in patients harboring L144P mutation. Our in-silico prediction and in vitro validation demonstrated that L144P mutation affects the channel function. We observed high on-target efficiency in the CBEs treated L144P mutant gene expressing HEK-293 cells. Strikingly, our evaluation of base editing efficacy using electrophysiology showed negligible channel function. We found that the editing bystander ‘Cs’ in the protospacer region led to a missense change (L143F) in evoCDA edited cells and only silent changes in BE4max edited cells. Upon investigation of the effect of the synonymous codon, our extended analysis revealed distortion of mRNA structure, altered half-life, and/or low abundance of the cognate tRNA. We propose that *KCNJ13*-L144P mutation or other genes that share similar genetic complexity may be challenging to correct with the current generation of CRISPR base editors, and a combinational therapy using CRISPR base editors with a tighter editing window and requisite cognate-tRNA supplementation could be an alternative therapeutic approach to restore Kir7.1 channel function in LCA16 patients. Other options for hard-to-rescue alleles could employ homology-directed repair using CRISPR/Cas9 nucleases, Prime editing, and AAV-mediated gene augmentation.

## Introduction

Point mutations in *KCNJ13* (MIM#603208) gene cause an autosomal recessive disease, Leber congenital amaurosis 16 (LCA16, MIM#614186)(1–4). Allelic heterogeneity in this gene is also observed in an autosomal dominant phenotype, Snowflake vitreoretinal degeneration (SVD, MIM#193230)(5). The LCA16 phenotype manifests in early childhood and is clinically diagnosed by pigmentary abnormalities in the retina, reduced or complete loss of electroretinogram (ERG) waveforms, nystagmus, photophobia, and progressive loss of central and peripheral vision leading to blindness(6, 7). *KCNJ13* gene encodes a *homo-tetrameric* inwardly rectifying potassium channel, Kir7.1, which is expressed in the apical processes of retinal pigmented epithelial (RPE) cells. The Kir7.1 channel is vital in RPE cells to maintain its membrane potential, ionic homeostasis in subretinal space, and phagocytosis of photoreceptors outer segment(8, 9).

There is no treatment for LCA16, but several approaches to rescue the Kir7.1 function as a therapeutic invention are in the trial. We previously reported an LCA16 patient with W53X mutation in the *KCNJ13* gene, which affected Kir7.1 channel expression and function(2). We also showed the rescue of channel function using translation readthrough inducing drugs (TRIDs) and successful gene augmentation therapy in an LCA16 patient-derived hiPSC-RPE model(9). Treatments using TRIDs are nonsense suppression specific, incorporating an amino acid to replace the stop codon. Depending on the near cognate amino acid inserted, the resulting protein may or may not lead to a functional channel like native Kir7.1.

In contrast, gene augmentation is a global approach for loss-of-function mutations. Still, the dominant-negative effect of mutation as seen in SVD and lentiviral and AAVs associated adverse immune responses make this approach questionable and underscore a need to develop new treatment options. Recently developed CRISPR/Cas9 genome editors offer a therapeutic opportunity(10–13). This approach involves the double-stranded DNA break formation followed by its repair either by donor-dependent homology-directed repair (HDR) or non-homologous end joining (NHEJ). The NHEJ forms on-target and genome-wide unintended indels, limiting its use as a therapy(14–18). Traditional CRISPR/Cas9 gene editing is more challenging for autosomal recessive cases of channelopathies requiring bi-allelic on-target edits to produce homo or hetero-multimeric channels. But if both alleles are edited differently, one with the desired change and the other with undesired substitutions or indels, the outcome might compromise channel function.

Most of the inherited ocular channelopathies are caused by point mutations(4). Therefore, CRISPR base editing can reverse the effect of mutation by changing a single base without the need for HDR or NHEJ. The LCA16 and SVD causing *KCNJ13* mutations are single nucleotide changes, and therefore, CRISPR-base editing via adenosine base editors (ABEs) for G>A (or C>T) mutations [c.158G>A; W53X, c.458C>T; T153I, c.496C>T; R166X, c.655C>T; Q219X, c.484C>T; R162W] or cytosine base editors (CBEs) for T>C (or A>G) mutations [c.359T>C; I120T, c.722T>C; L241P] in *KCNJ13* can conceivably rescue the channel function(1-3, 5, 19). As a proof-of-concept study, in the current report, we explored the activity of two CBEs (BE4max-WTCas9 and evoCDA-SpCas9-NG) at the recently reported L144P (c.431T>C) mutation site. The L144P mutation was selected as its role in LCA16 has not been established. Proline is a well-known secondary structure breaker(20) due to its structural rigidity, which complicates the wild-type ordered structure and might have a direct impact on the channel. In this study, two LCA16-patients harboring L144P mutations were clinically evaluated for the progression of the disease phenotype. To understand the possible molecular etiology due to L144P, we used multiple in silico tools to predict the mechanism of Kir7.1 channel dysfunction. We developed a stable HEK293 cell line as a heterologous overexpression system to validate the in-silico findings. We also tested the potential of CRISPR base editing to correct L144P mutation in our stable cell model. Although CRISPR base editing rescue retinal and visual function in LCA-RPE65 mice using adenosine base editors delivered by lentivirus,(20) Leu to Pro mutation is unique to be edited by CBEs as all the Pro-codon has two Cs which are not at the wobble position. Editing of two Cs using CBEs may lead to a preferred Leu (with or without synonymous variation) or undesirable Phe amino acid at 144. We evaluated two previously reported CBEs’ potential and a specific sgRNA targeting L144P mutation. Further, the Kir7.1 channel function was evaluated in edited cells with bystander silent variations by electrophysiology to elucidate the role of synonymous changes. We also examined the role of these synonymous variations on Kir7.1 mRNA stability and its translation by checking the abundance of tRNA for the Leu codon. The present study supports the need to translate the genotype data, exceptionally with silent variations, for use in the clinic.

## Results

### 1. Patients present clinical features of LCA16

We report a consanguineous family(21) (Family ID 6) of Arab descent with two affected female siblings, both diagnosed in infancy but now age 9 (patient ID 6-1) and 5 (patient ID 6-2) years. Both siblings had a characteristic *KCNJ13*-LCA16 phenotype with nummular pigment areas at the RPE level, especially over the posterior pole, macular atrophy, and optic disc pallor with retinal vessel attenuation (Fig 1). There were no signs of retinovascular changes or neovascularization as previously reported(22) Patient 6-1 had rotatory nystagmus with a best-corrected visual acuity (BCVA) of 1.48 LogMAR in the right eye and 1.6 LogMAR in the left eye. Her refraction was +1.50/-2.00×170 in the right eye and left eye +0.50/-2.25×10. Patient 6-2 has horizontal nystagmus and a left intermittent exotropia, and her BCVA was 0.80 LogMar in both eyes with a myopic astigmatism right eye −5.00/-2.75 x179 and left eye −5.00/-2.50×175. RETeval® flash and flicker ERG showed no detectable responses. SD-OCT revealed extensive loss of the ellipsoid zone in both patients along with retinal thinning and disorganization. The affected siblings were found to have a homozygous missense variant (c.431T>C, p.Leu144Pro) in *KCNJ13*, using Sanger sequencing(21). Parents did not require segregation as the variant was homozygous in both affected sisters. No disease-causing mutations were found in any other known retinal disease genes.

**Figure 1:**
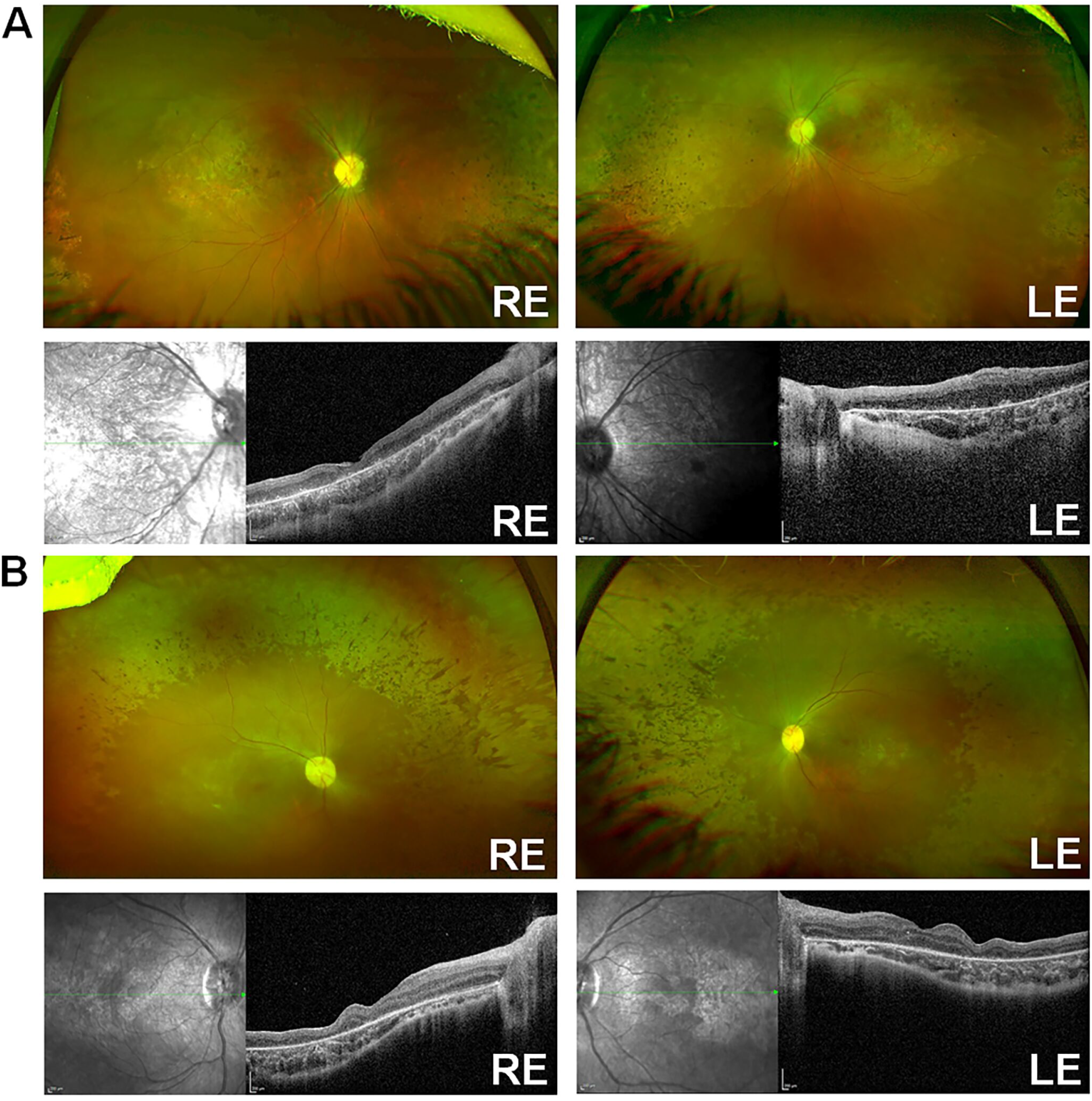
Clinical characterization of LCA16 patient with L144P mutation. [A] Optos and SD-OCT for Patient 6-1, [B] Optos and SD-OCT for Patient 6-2.

### 2. In silico tools predicted the L144P mutation as pathogenic

Amino acid L144 is located within the transmembrane domain-2 (TM-2) of the Kir7.1 protein [Fig 2a], which is important for channel gating and inward rectification. Although the leucine (non-polar) to proline (non-polar) change does not alter the charge of the TM-2 region, proline substitution can interrupt transmembrane alpha-helices and disrupt the protein structure. We used several computational tools to predict if the L144P mutation has any structural and functional impact on the Kir7.1 channel. All the seven computational algorithms (SIFT, PolyPhen-2, PANTHER, SNPs&GO, PROVEAN, PredictSNP, SNPA-2) used in this study identified L144P mutation as deleterious or disease-causing [Fig 2b]. This variant was found twice in a heterozygous state in 125,455 people (allele frequency = 0.0007971%) from multiple origins (gnomAD). I-Mutant tool predicted decreased stability of Kir7.1 protein due to L144P mutation in terms of reliability index (RI=4) and difference in free energy change values (L144P-WT=-1.30 Kcal/mol). The L144 amino acid residue of native Kir7.1 was highly conserved across multiple species, suggesting the Leu residue’s functional relevance at the specific position [Fig 2c]. A SNAP-2-generated heatmap provided the possible substitution at each position of Kir7.1 protein, where a score >50, shown in red, indicated a strong signal for pathogenicity of L144P (Score=62, 80% accuracy) [Fig 2d]. The protean-3D subroutine in DNASTAR showed minor differences in the secondary structure of Kir7.1 (Dihedral angles, φ;-62.8°, ψ; −36.5°, ω;173.8°) due to L144P substitution (Dihedral angles, φ;-67.1°, ψ; −34.6°, ω;170.7°) [Fig 2e]. Similarly, SOPMA tool reflected scarcely any differences in the overall composition of Kir7.1 secondary structure including alpha helix (WT=30.56%, L144P=29.44%), beta-turn (WT=4.72%, L144P=5.28%), extended strand (WT=23.33%, L144P=22.50%) and random coil (WT=41.39%, L144P=42.76%). However, the L144P mutation-induced remarkable variability (marked in black rectangles) in the basic secondary structure at the C-terminal of Kir7.1 [Fig 2f]. The C-terminus is the critical determinant for the membrane localization of Kir7.1 and contains the putative phosphorylation sites S287 (Protein Kinase A), T321, and T337(Casein kinase II)(23). Alanine substitution of these phosphorylation sites did not affect the Kir7.1 trafficking to the membrane. But changes in the beta-turn and alpha-helix proportion at these phosphorylation sites and proximal and distal C-terminus could affect the cytoplasmic pore formation in the tetrameric channel and inwardly rectification(24–26).

**Figure 2:**
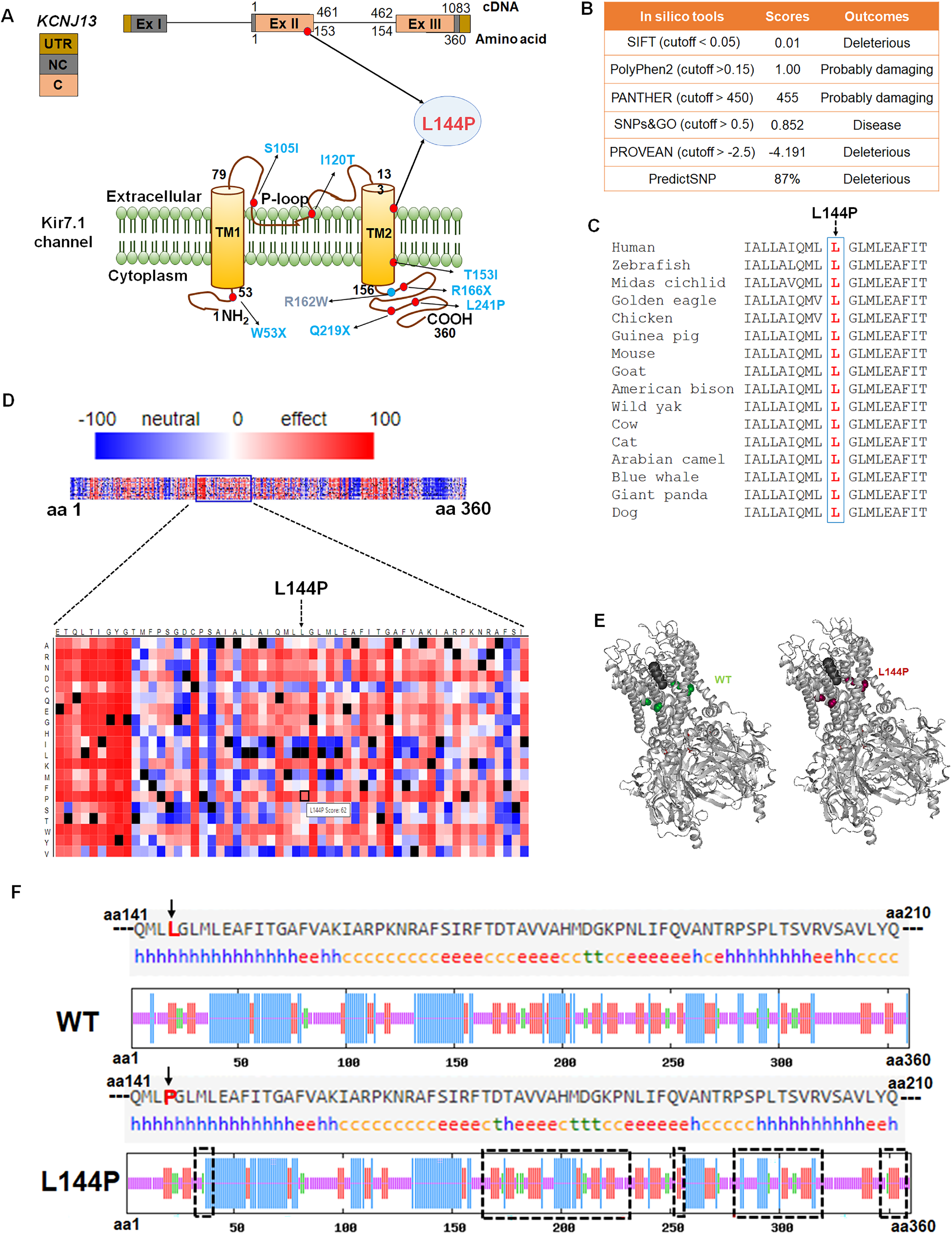
Pathogenicity of the L144P mutation. [A] Genomic and protein location of L144P mutation. [B] In silico tools to predict the pathogenicity of L144P substitution. [C] Conserved L144 amino acid. [D] Heatmap of Kir7.1 substitution predicted by SNAP-2 of L144P pathogenicity. [E] DNASTAR (protean-3D) generated secondary structure of native and L144P Kir7.1. [F] SOPMA tool reflecting the proportion of alpha-helix (blue ‘h’), extended strand (red ‘e’), beta-turn (green ‘t’), and random coil (orange ‘c’) in native and L144P Kir7.1. The black rectangle shows the Kir7.1 region with the variable arrangement of basic secondary structure induced at the C terminal of Kir7.1 due to L144P substitution. (Figure 2_Source Data 1 contains full images generated by SOPMA tool).

### 3. L144P substitution altered the protein localization

To determine whether the L144P substitution alters the localization of the Kir7.1 channel, we examined the cellular distribution of native and L144P Kir7.1 in HEK293 cells. As reported earlier, the cell membrane showed signs of native Kir7.1 expression [Fig 3a](27). However, cells transfected with mutant eGFP-L144P-Kir7.1 plasmid showed fluorescence signal in the cytoplasm and other organelles, indicating that L144P substitution affects the normal regular protein transport via classical conditioning ER/Golgi pathway [Fig 3b]. Staining with an ER tracker showed that the mutant protein was primarily distributed in the endoplasmic reticulum [Fig 3c]. The loss of membrane expression of these mutant proteins could be due to a defect in its ER exit.

**Figure 3:**
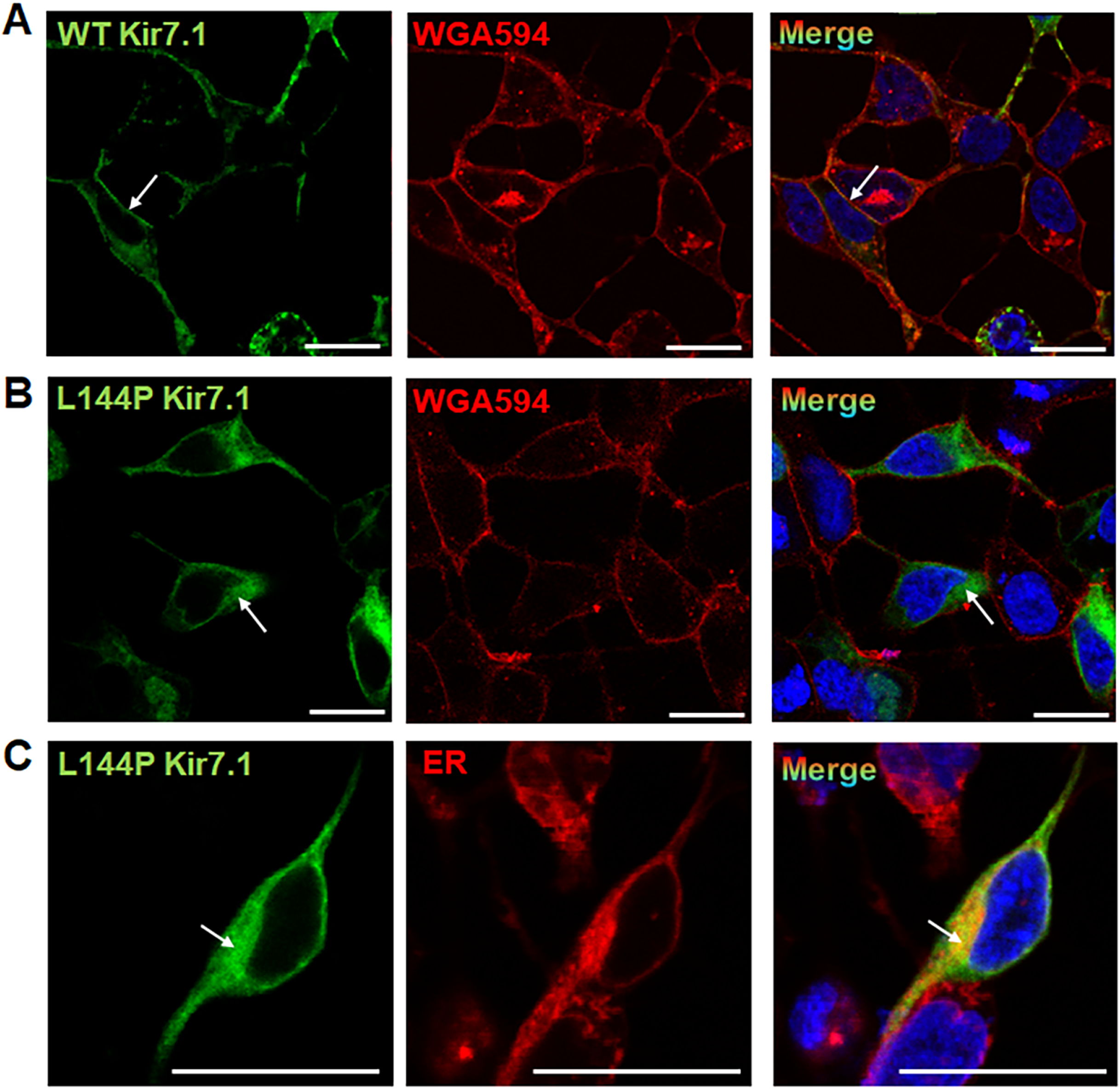
Localization of Kir7.1 protein in HEK293 cells. The cells were transfected with GFP-WT-Kir7.1 or GFP-L144P-Kir7.1 plasmid. [A] Native Kir7.1 (green) expression in the membrane (red). [B] Mutant L144P-Kir7.1 (green) expression in the cytoplasm and other organelles. [C] Localization of a significant proportion of L144P-Kir7.1 in the Endoplasmic Reticulum (red). White arrows show the colocalization of Kir7.1 with membrane, cytoplasm, or ER. Scale: 25 μm. (Figure 3_Source Data1 contains more images from different fields).

### 4. CRISPR Base editing of L144P mutation resulted in gene correction with synonymous changes

Base editors can be used to precisely introduce transition edits and reverse transition point mutations. We wanted to test if this approach can ameliorate the mutant phenotype of Kir7.1-L1444P gene function by directly correcting the C>T mutation. To install this edit, we first designed sgRNAs to the target L144P mutation in exon 2 of *KCNJ13* for two cytosine base editor mRNAs, BE4max-WTCas9 and evoCDA-SpCas9-NG. CBE mRNAs chemically modify the targeted C to T (or G> A on the opposite strand) by engaging its deaminase domain with WT Cas9 or NG-SpCas9 and a 20-bp sgRNA. The WT Cas9 best recognizes the NGG PAMs and has low activity on NGA or NAG PAMs. SpCas9-NG recognizes NG PAMs but has some activity on NA PAMs and seems to do best on NGNG PAMs(28, 29). The activity window of BE4max CBE mRNA ranges from positions 4 to 9, but some action was also observed at 3 and 12 locations in the protospacer sequence, counting the PAM position as 21-23(30). The evoCDA CBE mRNA activity window ranges from positions 1 to 13, but the peak editing has been observed at positions 4 to 6 of the protospacer (31). All three gRNAs are predicted to have some off-target activity against the human genome in silico (Supplementary Figure 2). The sgRNA-2 had the highest on-target activity at the desired location (Cytosine-6; C6), which is a prerequisite for efficient biallelic correction and, therefore, was chosen for CRISPR base correction L144P.

The CBE mRNA (BE4max-WTCas9 or evoCDA-SpCas9-NG) and the chemically modified sgRNA-2 were nucleofected into L144P mutant stable cells at a 3:1 ratio by weight. The sgRNA-2 guide sequence spans a region containing multiple bystander ‘’C’s, and within the sgRNA, these ‘C’s are labeled as C2-C17. [Fig 4a]. First, we tested the % correction of pathogenic L144P missense mutation in the treated cells by genomic analysis. Reverse transcription PCR of mRNA from the evoCDA treated cells and deep sequencing analysis showed higher editing of the desired C6 (78.8% ±7.00) base than BE4max CBE treated ones (66.27% ±4.25) [Fig 4b]. We also examined edits at bystander Cs in the protospacer region. Cells edited with either editor showed editing of bystander Cs (C2-C5, C10-C17) within the protospacer region [Fig 4c]. BE4max mRNA-treated cells showed a narrower editing window with only silent bystander mutations and most of the reads with Leu at 144 (59.0% ± 6.1) [Fig 4c]. In contrast, evoCDA mRNA showed missense bystander mutations across a wide editing window, and only ∼ 2% of reads were with Leu at 144 [Fig 4c, 4d & 4e]. The editing of C2 (non-silent) and C11 (non-silent) to T led to a missense mutation phenylalanine (F) at amino acid 143 (61.4%) and 146 (4.14%) in evoCDA treated cells. At the same time, these Cs remain untouched in BE4max-treated cells. None of the CBE-treated samples showed higher editing activity outside the protospacer. In addition, we also examined the on-target indel formation by CBEs, which showed variability in three independent experiments. BE4max CBE treated cells had a higher indel frequency (13.8% ±4.8) compared to evoCDA CBE ones (8.3% ±5.9) [Fig 4c]. Untreated cells (UT) were used as a reference (Supplementary Figure 3). The observed efficient editing prompted us to pursue the functional evaluation of edited cells.

**Figure 4:**
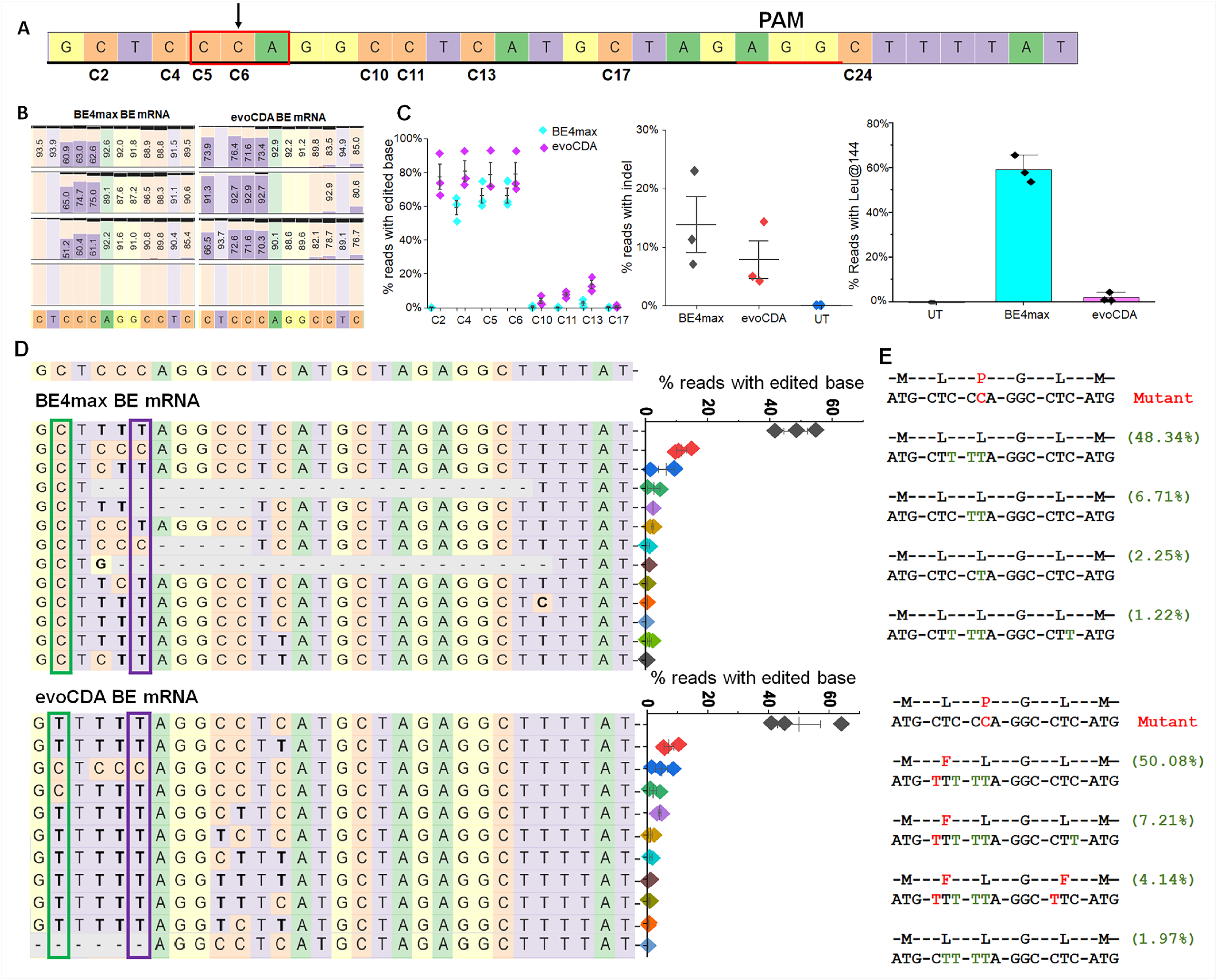
Efficiency of CBE mRNAs (evoCDA and BE4max) in editing the L144P mutant allele and on-target analysis in CRISPR base-edited cells. [A] Distribution of C (C2 to C24) around the target C6 location. The sgRNA sequence is underlined by black and the PAM by red. Nucleotides are marked by unique colors (A; green, G; yellow, C; orange, T; purple. [B] Frequency of nucleotides around sgRNA GCTCCCAGGCCTCATGCTAG location as observed in sequencing reads from treated cells. The black horizontal bars indicate the % of reads for which that nucleotide was deleted [C] % Editing of the target (C6) and bystander (C2-C5, C10-C17) ‘C’ to ‘T’ and % indels by BE4max and evoCDA mRNA as observed in three independent experiments. The error bars represent the SE. [D] Top 10-13 reads showing the nucleotide distribution around the cleavage site for sgRNA. A dashed line ‘-’ designates the deletion of bases while substitutions are shown in bold. Reads generated by BE4max mRNA treatment show C2 location untouched and C>T conversion at the desired location. A scatter plot shows the frequency of each read observed in treated cells (n=3). A green rectangle box marks the C2 (aa-142) location and the purple C6 (aa-144). The lower panel shows the reads generated by evoCDA mRNA treatment and their frequency. C2 is edited to T in most of the reads, giving rise to a missense mutation phenylalanine (F) at amino acids 143 and 146. [E] Amino acid conversion at the respective location for the 4 top reads (based on frequency) shows the synonymous (green) and missense (red) variants generated due to bystander C edits. Figures presenting pooled data are represented as mean ± SEM. (Figure 4_Source Data 1-3 contains raw and analyzed NGS files for BE4max treated, evoCDA treated and untreated (n=3) samples).

### 5. CRISPR base editing of L144P showed protein restoration in the membrane

As synonymous variations are assumed not to alter the protein function, we tested our CRISPR-edited pool of cells to restore Kir7.1 protein levels and functions. Our immunocytochemistry assay using primary antibody against the GFP showed cell membrane localization of Kir7.1-GFP protein for some of the edited cells in the analyzed BE4max treated pool [Fig 5a], akin to the localization of wild type Kir7.1 [Supplementary Figure 1]. Kir7.1 within the evoCDA treated cell pool showed the membrane localization despite the L143F mutation. Both the edited lines showed protein restoration in some of the cells; these were further evaluated to compare the biophysical properties of the Kir7.1 channel.

**Figure 5:**
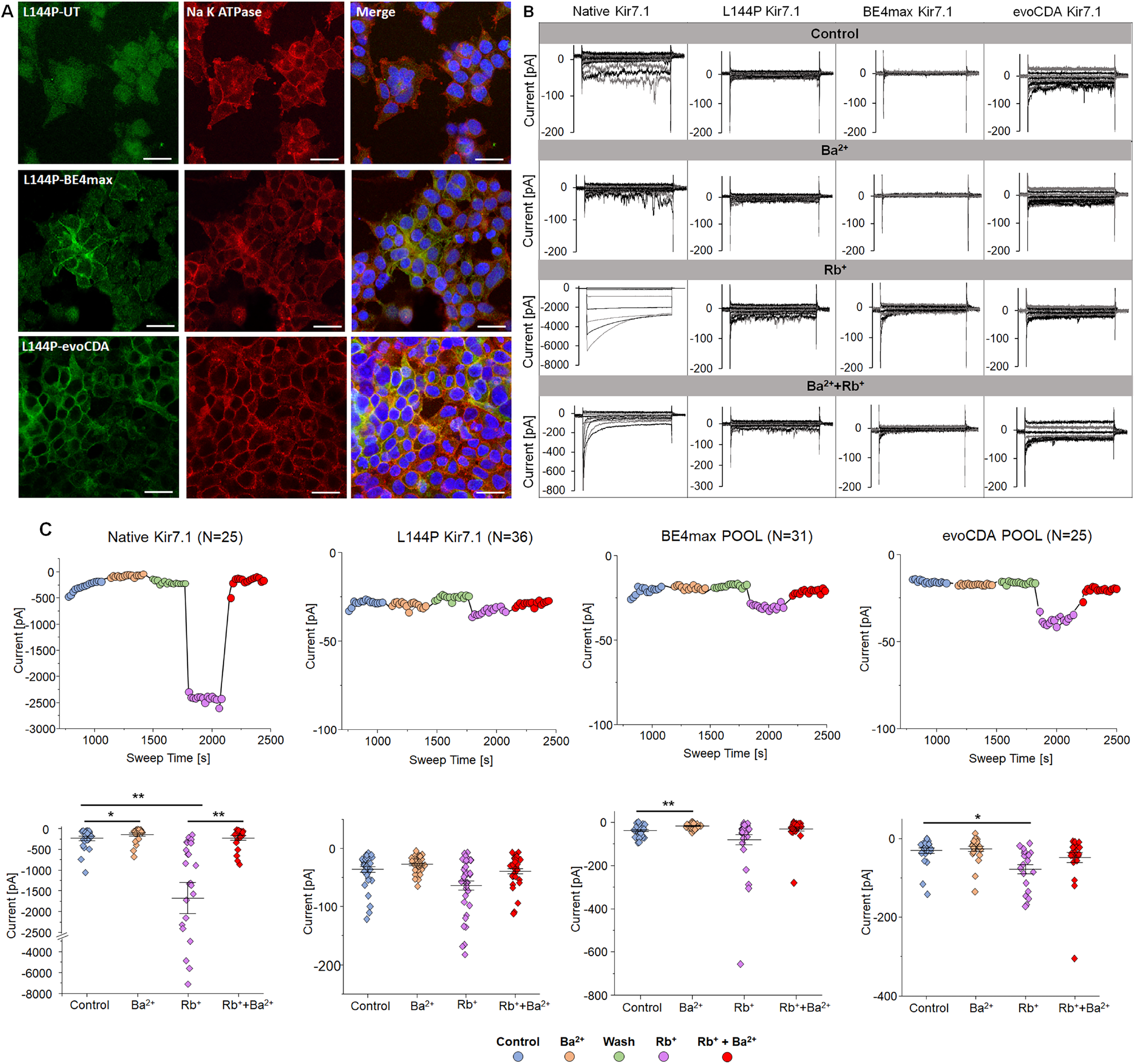
Protein localization and K^+^ current after L144P base editing. [A] Kir7.1 Protein localization in L144P untreated and base edited cells assessed by immunocytochemistry. Scale: 25 μm. [B] K^+^ current in WT, L144P-mutant, and base edited cells. Currents were elicited by 550 ms voltage steps from −150 mV to +40mV (20mV increments) followed by a step to −10 mV (250ms). [C] Current-Sweep Time plot from a representative cell in physiological external solution (blue), external solution with 10 mM Ba^2+^ (yellow), wash with external solution (green), external solution with 140 mM Rb^+^ (purple) followed by external solution with Rb^+^+Ba^2+^ (red). The average current profile for WT, mutant, or base edited pools (n=25 to 36) are shown in the presence of respective solutions. [*p< 0.05, **p< 0.001]. Figures presenting pooled data are represented as mean ± SEM. (Figure 5_Source Data 1 contains the raw files generated from automated patch clamp (APC) system without excluding any data).

Whole-cell current was measured in 5mM [K+] in L144P mutant and based on edited pooled cells compared to WT cells to study channel functional expression. The cells were exposed to a 550 ms voltage pulse from −150 mV to +40mV (20mV increments) from a holding potential of −10 mV. The step current-voltage plot and the current-sweep time plot for a representative cell type are shown in different treatment solutions in Figures 5b & 5c. Our electrophysiological outcome showed salient features of Kir7.1 current in physiological solution in WT cells. We observed a weak inward rectification and increase of current at negative potentials in WT cells which are inactivated to near zero at more depolarized voltages. The WT FRT stable cells produced an average Kir7.1 current of −0.23± 0.05 nA (n=24) at −150 mV in 5mM K^+^ [Fig 5c]. The WT channel also showed an increased current in external Rb^+^ and inhibition in the presence of Ba^2+,^ as expected for a normally functioning K^+^ channel. The Ba2+ sensitivity for the Kir7.1 channel was low but resulted in a 2-fold decreased current (−0.11± 0.03 nA, *p*; 0.04). On an average, a ∼7-fold increased conductance (−1.67± 0.37 nA, *p*; 0.0008) was observed following the addition of 140 mM Rb^+^ to the external solution while addition of 20mM Ba^2+^ in the Rb^+^-external solution caused a 7-fold decrease in the Rb^+^ signal (−0.22± 0.05 nA, *p*; 0.0008).

The L144P single-cell patch-clamp (n=36) showed significantly lower current amplitude (−0.04 ± 0.004 nA, *p*;0.002) than the WT cells in 5mM K^+^ but negligible response to external Rb^+^ (−0.05 ± 0.006 nA, *p*;0.05), Ba^2+^ (−0.03 ± 0.002 nA, *p*;0.098) and Rb^+^+Ba^2+^ (−0.04 ± 0.004 nA, *p*;0.902). Surprisingly, we did not observe the restoration of K^+^ current in a pool of BE4max edited cells (n=27, −0.04 ± 0.004 nA, *P*;0.862). Most of the cells had synonymous variation, and current amplitudes were comparable to mutant L144P channel expression. These cells showed a significant block of K^+^ current in response to external Ba^2+^ (−0.02 ± 0.002 nA, *P*;0.001) but moderate increase in the current amplitude with respect to Rb^+^ (−0.04 ± 0.004 nA, *P*;0.755) [Fig 5b & 5c]. In the pool of BE4max edited cells, only one cell (3.6%) showed a K^+^ conductance (−0.07 nA) and Rb^+^ (−0.31 nA), and Ba^2+^ response (−0.01 nA), which may be a rare cell containing in-frame indels or the WT genotype and no synonymous variations. This could also result from a hetero-tetrameric channel expression in the cell. The stoichiometric ratio of the WT channel is equal/more than the one with synonymous changes. The evoCDA treated pool of cells also showed a response similar to mutant cells due to the L143F and synonymous variations. There was no difference in the average current amplitudes measured at −150 mV in the presence of 5 mM K^+^ (−0.03±0.01 nA) and Ba^2+^ (−0.03±0.01 nA). Unlike native Kir7.1, the average increase in the current amplitude was only 2-fold in the presence of Rb^+^ (−0.07±0.01 nA, *p*; 0.01). These results suggest that there may have been a significant loss of WT channel function due to silent variations.

### 6. Genomic off-target effects of L144P targeting sgRNA were fewer with BE4max CBE as compared to evoCDA CBE mRNA

Screening the computationally predicted potential off-target (OT) sites within the human genome is necessary to evaluate the safety and efficacy of CRISPR-based therapies. An in*-*silico search analysis via Cas-OFFinder(32) found 1136 OT sites [Fig 6a], each having 1-3 mismatches concerning sgRNA-2, with or without a DNA/RNA bulge of 1 nucleotide. Most of the identified sites were with 3 mismatches and a single RNA bulge (n=790), followed by 3 mismatches with a DNA bulge (n=266) [Fig 6a & b]. We tested the efficiency of modifying selected OT sites [Fig 6c] to two CBEs in treated L144P stable cells. Consistent with other published studies, our study showed that CBEs could induce genome-wide unintended genomic modifications. Differences in their deaminase properties can affect their editing profile and efficiency(33–36). Deep sequencing analysis of 12 putative off-targets showed that BE4max had high activity (30.75% substitution, 2.70% indel at OT3) at only one of the genomic locations. The reads from untreated cells showed a baseline indel formation of ∼0.1%, and we set a threshold of 0.033% for our off-target assay based on the base level substitutions and indels in reference cells. The evoCDA CBE also had high OT activity at OT3 and OT6, and OT7 sites. Overall, evoCDA CBE had comparatively higher substitution and indel frequency at all OT sites with detectable modifications in this assay. The OT3 is in the intronic region of *PDZD4* (Supplementary Table 4), >13 K-bp away from the splice site. Therefore, variants at this OT site are unlikely to impact the PDZD4 or Kir7.1 protein function. BE4max did not show OT activity at other sites higher than the control treatment. At the same time, evoCDA CBE had a more comprehensive range of substitution frequency (∼0.2-51%) and some indel formation (∼0.1-4.2%) across all the remaining sites [Fig 6d].

**Figure 6:**
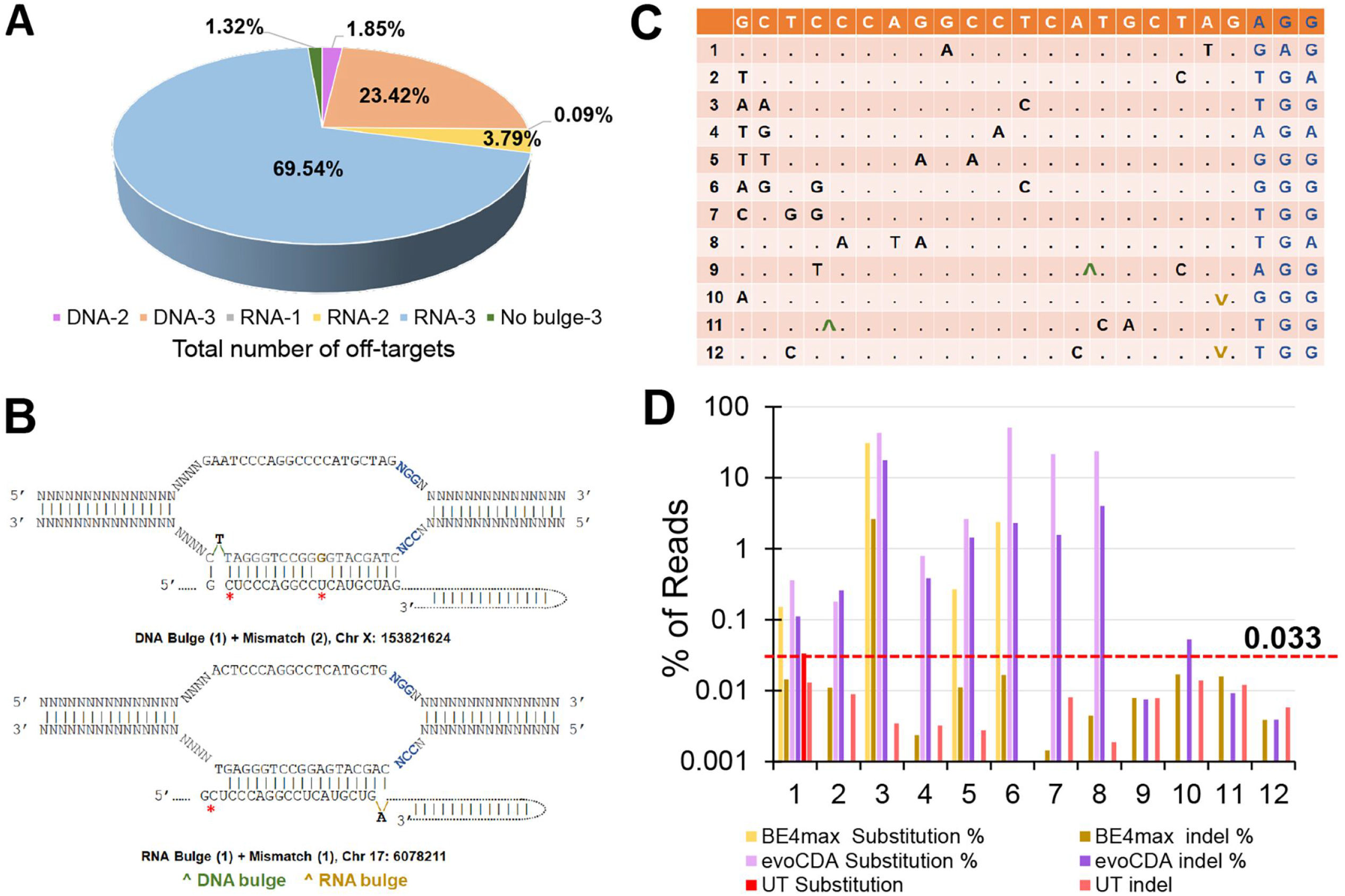
Off-target analysis of L144P sgRNA in CRISPR base-edited cells. [A] Total number of off-targets, each having 1-3 mismatches with or without a single RNA/DNA bulge as observed in in-silico analysis. (Figure 6_Source Data 1 contains the complete list of off-targets). A large fraction of OT sites was with 3 mismatches and a single RNA bulge, followed by 3 mismatches and a single DNA bulge. [’DNA-2’ in the pie chart represents the single DNA bulge with 2 mismatches, ‘RNA-3’ represents the single RNA bulge with 3 mismatches, and so on]. [B] Representation of DNA and RNA bulge with 1 or 2 mismatches with respective L144P sgRNA-2. [C] The 12 potential off-target sites with mismatches and DNA/RNA bulge and PAM site were screened by deep sequencing. [D] % substitution and indel frequency of BE4max and evoCDA CBEs at 12 off-target sites (Figure 6_Source Data 2 contains the NGS files in fastaq.gz format). L144P cells sham-nucleofected were used as reference. A threshold (red dashed line) was set at 0.033 based on the base level substitutions and indels in reference cells.

### 7. The synonymous variation is a by-product of editing that alters the mRNA stability and, thereby, protein synthesis

Synonymous changes are often assumed to have minimal effect on gene/protein function. However, several studies have implied that synonymous variations could disrupt transcription, splicing, mRNA stability, and translation kinetics(37–39). Because we did not observe robust rescue of Kir7.1 channel function in our base edited cells, we further examined the role-play of synonymous variation observed due to bystander ‘C’ editing in altering the Kir7.1 channel function. The BE4max CBE edited cells were primarily edited at the target nucleotide and nearby sites yielding only synonymous outcomes. Therefore, only these cells were flow-sorted to select single-cell clones (Figure 7_Source Data 1 contains the flow sorting images). Most of the sequence-verified clones (n=18) were either CTC [L143]-TTA [L144] (L144bystander-clone I, 27.8%) or CTT [L143]-TTA [L144] (L143-L144bystander-clone II, 38.9%) [Fig 7a], consistent with our pooled sequencing results. The rest of the clones were either unedited or contained indels in *KCNJ13*. The clonal cells of types I and II were further propagated to study the protein localization and rescue of Kir7.1 function. An immunocytochemistry assay demonstrated that a fraction of protein is trafficked to the membrane in both clonal types. However, a large proportion is still confined to cell cytoplasm and organelles [Fig 7b]. This result suggested that these silent variations might alter protein folding, hindering oligomerization to form a tetrameric channel, thus failing during ER quality control.

**Figure 7:**
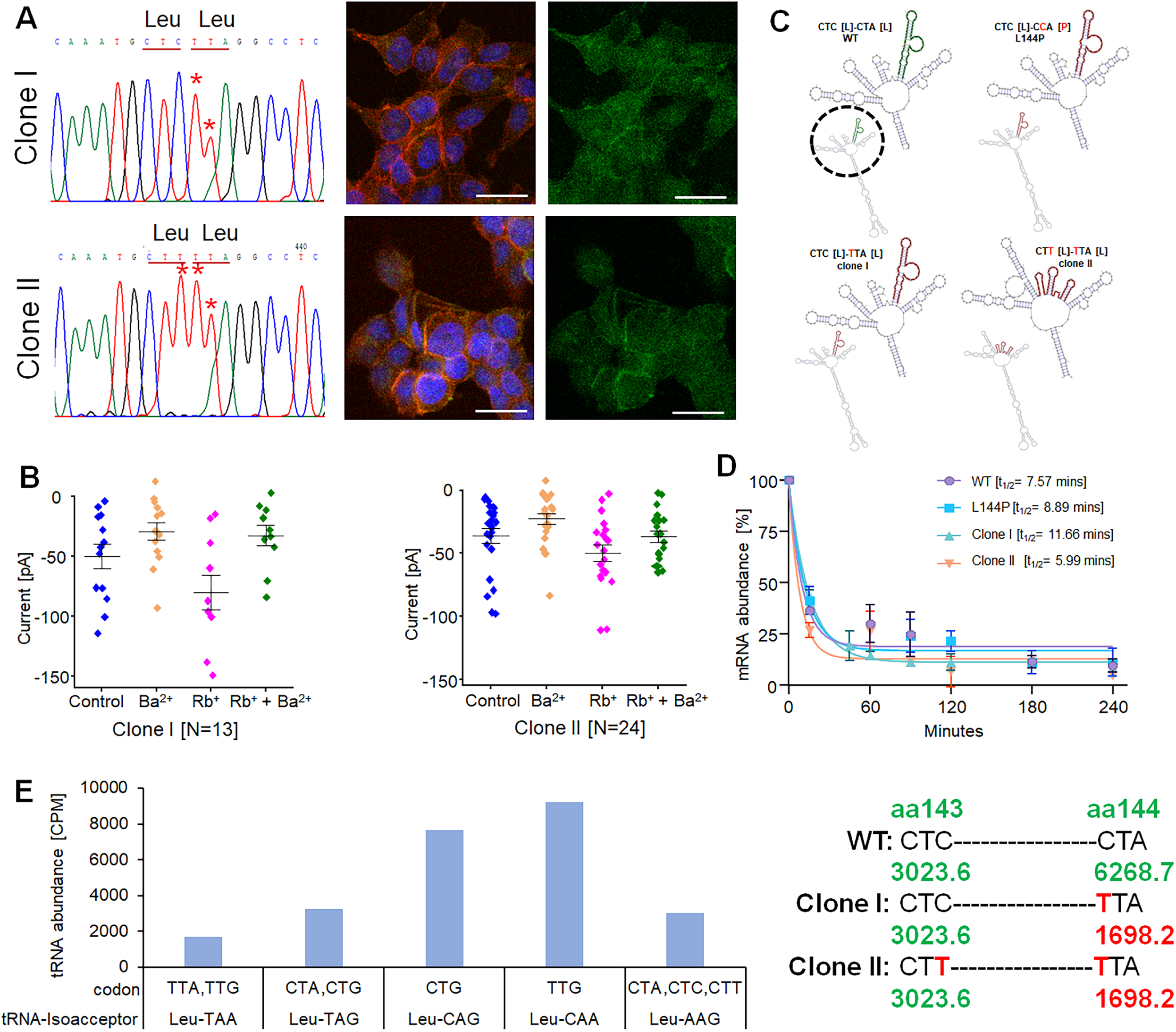
Role of synonymous variations observed as a by-product of CRISPR base editing. [A] Chromatogram of two single-cell edited clones (clone I and clone II) and the respective Kir7.1 expression. (Figure 7_Source Data 1 contains the flow sorting images and Source Data 2 contains chormatograms from other edited and unedited cells, n=3). [B] K^+^ current profile of two single-cell clones with a synonymous variation. (Figure 7_Source Data 3 contains raw files generated from APC). [C] The predicted optimal mRNA secondary structure of global sequence from WT, L144P mutant, and two edited clones. The enlarged mRNA structure within the black circle highlights the changes in the disruptive region of the sequence. [D] Half-life of mRNA after ActD treatment. [E] Leu-tRNA abundance in HEK293 FRT stable cells. The y-axis represents the sample’s CPM [counts per million] of tRNA. The x-axis represents the codons with their respective tRNA-anticodons. The right panel shows Leu’s abundance (CPM) at 143 and 144aa locations in WT, clone I, and clone II cells. Figures presenting pooled data are represented as mean ± SEM. (Figure 7_Source Data 4 contains the tRNA sequencing data from HEK293 FRT stable cells for Leu codon).

In patch-clamp experiments on the CRISPR edited cell clones, clonal type I (14 clones) and II (24 clones) showed that these cells do not have K^+^ conductance like native Kir7.1. Also, we did not observe any changes in the current amplitude mediated by the addition of Ba^2+^, Rb^+,^ or Rb^+^+Ba^2+^. Although the K^+^ current amplitude was too low (−0.06± 0.01 nA) in clone I cells, a 2-fold decreased Ba^2+^ current (−0.03± 0.007 nA, *p*;0.06) and a 2-fold potentiated Rb^+^ response (−0.13± 0.03 nA, *p*;0.05) was observed. The type II clones showed the current amplitudes close to uncorrected L144P cells. We observed −0.04± 0.006 nA K^+^ current, −0.05± 0.006 nA Rb^+^ Current and −0.02± 0.004 nA Ba^2+^ current in these cells [Fig 7b].

Since we did not observe the robust K+ conductance in the edited clones, we evaluated if these synonymous variations impact mRNA structure or stability. To understand the effect of these silent variations on mRNA structure, we used an in-silico tool RNAsnp, which can predict the changes in mRNA secondary induced by SNPs based on global folding(40). The Kir7.1 mRNA sequence and the two silent variations were given input in the RNAsnp server using Mode 1. The tool predicted that the base-pairing probabilities in WT, L144P, and type I clones were similar, and the mutation (CTA>CCA, L144P) or C>T silent change (CTA>TTA, L144L) did not have any impact on the mRNA structure. However, the other synonymous variation in type II clones (CTC>CTT, L143L) disrupted the mRNA structure in the region overlapped with the position of C>T variations (L143 and L144) [Fig 7c]. These results indicated that structural disruption of mRNA might potentially affect the mRNA stability and translation of the total number of transcripts at a time. To determine if the mRNA stability is affected in a codon-dependent manner, we treated the cells with Actinomycin D to block the transcription of new mRNA. The decay of the existing mRNA was measured by performing a time-course assay [0-4 hours] to check the mRNA abundance. Interestingly, the half-life of Kir7.1 mRNA in type II clones was similar to that of the WT mRNA, while mRNA in type I clones showed a comparatively higher half-life than the WT Kir7.1 transcript [Fig 7d]. Altogether, these results suggest that clone I mRNA may accumulate in cells due to its higher half-life, and structural disruption in clone II mRNA might impact the gene expression by altering the translation of Kir7.1 mRNA into protein.

The other factor that could alter the translation kinetics is tRNA abundance for a specific codon. Therefore, we determined if there is any difference in tRNAs abundance for the Leu codon (n=6, CTC, CTA, CTG, TTA, TTG, CTT) in the cells, as codon biases and differences in the availability of cognate tRNAs may alter the translation rate and the subsequent protein co-folding mechanism. Our tRNA sequencing from HEK293 FRT stable cells using an Illumina NextSeq platform showed comparatively reduced availability (counts per million, CPM) of a cognate-tRNA for ‘Leu’ (TTA), which was generated due to bystander ‘C’ edits [Fig 7e].

## Discussion

Two pediatric siblings were previously included in a brief study showing a homozygous missense variant in *KCNJ13*; c.431T>C, p.(Leu144Pro) (L144P)(21). Herein, we report their clinical phenotype consistent with other LCA16 patients harboring different *KCNJ13* point mutations (1–3). Both patients were noted to have nystagmus, early-onset loss of vision, pigmentary retinopathy with degeneration of the ellipsoid zone, attenuation of retinal vessels, and altered electroretinogram. There were no signs of retinal neovascularization or fibrovascular changes, as seen in patients with the p.Thr153Ile mutation, although the siblings are still relatively young(22). The L144P mutation is in exon 2 of *KCNJ13*, which forms the TM-2 domain of the Kir7.1 channel. Our prediction based on different in-silico tools identified this mutation as pathogenic, with altered stability. The mutation introduced salient variability in the basic secondary structure at the C-terminal of Kir7.1 protein, which contains the signal for membrane localization. This prediction was further supported by our findings in a heterologous expression system, where we found that L144P mutation causes a defect in its trafficking and leads to impaired membrane expression. The mutant protein was accumulated in the cytoplasm, ER, and other organelles, indicating a defect in the folding/assembly of tetrameric subunits or post-translational modifications. Our electrophysiology assay in HEK293 stable cells demonstrated that this mutation leads to a non-functional channel with a compromised inward rectification. Extracellular Rb^+,^ which amplifies the inward K^+^ current in a native Kir7.1 channel, Ba^2+^, which blocks the channel activity, did not change the current amplitude in L144P expressing stable cells.

Base editing is a powerful tool to correct point mutations, but membrane proteins like Kir7.1 are complicated to repair and restore function because of their multimeric structure. In addition, proline codons (CCU, CCC, CCA, CCG) with two adjacent Cs are challenging to correct without bystander edits. Cytosine base editing of proline codons may generate Ser, Leu, or Phe, one of which may lead to the desired edit back to a wild type state, while others would lead to a missense mutation and might have detrimental protein function. Moreover, the L144P site has multiple Cs within the activity window of CBEs, similar to other disease-associated alleles like APOE4 (Alzheimer’s disease)(41) and HBB (β-thalassemia)(42), which can potentially cause deleterious effects. The L144P mutation renders the Kir7.1 channel non-functional, and we attempted to rescue its function through CRISPR base editing.

In this study, we demonstrated the activity of two cytosine base editors, BE4max-WTCas9 and evoCDA-SpCas9-NG, to edit proline to leucine at 144. Our HEK293 stable cell lines expressing the mutant and control WT channel were precious to quickly test this approach since we did not have access to the patient’s iPSC-RPE cell lines or a mouse model harboring this mutation. Using these stable lines, our study generated new knowledge of correcting a *KCNJ13*-L144P mutation as the gene sequence at this location is unique with multiple bystander Cs. Also, the Pro mutation at 144 changes the codon [CtA>CcA] with two Cytosine bases, which fall in the editing window of CBEs. Although the codon has two cytosine bases, the redundancy of the Leu codon [CtA or ttA] generates a WT Kir7.1 protein sequence when corrected by CBEs. The CBE mRNA’s electroporation with a sgRNA resulted in high-frequency editing of the target site (60-80%). Other base editing outcomes producing bystander ‘Cs’ in the protospacer region led to a missense change (L143F) in ∼61% of evoCDA edited cells, which affected the Kir7.1 function. In BE4max edited cells, most of the reads had silent variation (∼59%), while very few reads had a WT gene sequence (∼3%). Our on-target genomic analysis showed comparatively higher indel frequency in BE4max treated cells than evoCDA ones, likely due to excision of the uracil intermediate by cellular DNA repair machinery.

Although CBEs can have high editing frequency at the target location, the therapeutic benefits of L144P-CRISPR base correction depend on its efficient activity without generating any bystander synonymous variants. The bystander base edits around the intended on-target site of base editors may present challenges for CRISPR base editing strategies looking to correct mutant alleles. EDIT-101CRISPR/Cas9 clinical trial (NCT03872479) of the CEP290 gene in LCA10 patients anticipated that 10% editing of foveal cones would lead to near-normal vision(42). For correction of L144P and other mutations with this type of sequence complexity, CBEs can be evolved further to have a tighter editing window with limited indels to increase the 3% reads (as observed in BE4max edited cells with WT genotype) to a minimum of 10% for clinical outcomes.

Additionally, our off-target analysis of 12 predicted sites showed higher activity of CBEs at one of the loci, *PDZD4*. The gene has not been reported to express in RPE or other retinal layers. As the activity region was found in its intron, it might not harm PDZD4 or Kir7.1 protein function. As consistent with other reports, BE4max was more specific based on its minimal activity at other off-target sites relative to evoCDA. We anticipated that L144P is an extraordinarily hard allele to edit because of many bystander Cs around the target site and within the editing window of CBEs.

This study also evaluated the Kir7.1 channel expression and function in the edited cells. We observed that some of the edited pool cells had membrane localization of the Kir7.1 channel comparable to WT protein. From BE4max edited pool, which also carried silent changes, electrophysiological analysis of flow-sorted single-cell edited clones showed minimal channel function compared to the native Ki7.1 level. We ruled out that the L144P mutation could act as dominant-negative and impact the channel function in edited cells because the patients’ parents with heterozygous mutations were unaffected. One of the limitations of our study is we did not explore the role of indels on channel function. The stoichiometric ratio of Leu-edited and indel/or unedited alleles in a cell could also alter the protein function.

Protein expression is regulated by a highly structured coding sequence of mRNA through changes in its half-life(43). We observed distortion of mRNA structure in the region of L144P, and inferred that this could be a reason for their altered half-life and subsequent translation. We extended our findings by describing the effect of synonymous mutation on mRNA stability. Our tRNA sequencing in the WT stable cells showed the low cognate-tRNA abundance for the synonymous codon, which was observed as a by-product of CRISPR base editing. Notably, the ‘TTA’ codon resulting from the synonymous variation had the lowest abundance (CPM=1698.2) among all the Leu-tRNAs, suggesting that ‘TTA’ is not the preferred (’optimal’) synonymous codon and could result in ribosome-halting, altered elongation rate during translation, or chaperone-assisted cotranslational protein folding.

In summary, this study generated several significant new findings. We demonstrated the potency of CRISPR-mediated base editing to correct the L144P mutation. Still, We indicated that synonymous bystander edits and unintended on-target effects of the cytosine base editor significantly impacted gene function, especially for its multimeric protein assembly which requires bi-allelic on-target edits. Due to its sequence complexity, we anticipate that the L144P locus poses some challenges to correct with the current generation of CRISPR base editors. Our study emphasizes the need to perform functional studies with on-target edited alleles to confirm the therapeutic potential of base editing. Improving potential therapies using modified CBEs with tighter editing windows(36, 44), supplementation of cognate-tRNAs, or both. Prime editing could precisely correct the disease mutation without introducing bystander edits. While recent insights into the mechanism of prime editing have determined that the intentional introduction of silent bystander edits can improve editing efficiency(45), this study suggests that caution is warranted to avoid detrimental outcomes of silent editing. Further optimization of genome editing approaches or gene delivery of the Kir7.1 channel could prevent vision loss in L144P-LCA16 patients.

## Methods

### Clinical evaluation of LCA16 Patient harboring L144P mutation

The molecular testing and clinical characterization had local approval through Moorfields Eye Hospital and adhered to the tenets of the Declaration of Helsinki. Informed consent was obtained from all participating individuals. The ophthalmic evaluation included best-corrected visual acuity, orthoptic assessment, cycloplegic refraction, slit-lamp anterior segment, fundus examination with ultra-widefield color fundus imaging with the Optos, SD-OCT (Spectral Domain Optical Coherence Tomography), and electroretinography using the RETeval® as part of routine clinical care.

### 2. Molecular genetic testing to identify the mutation

Molecular genetic testing was performed on genomic DNA extracted from blood using retinal dystrophy targeted gene panel testing through the Rare & Inherited Disease Genomic Laboratory at Great Ormond Street Hospital (London, UK) or Blueprint Genetics (Helsinki, Finland). Coding exons and flanking intronic regions of genes, including *KCNJ13* [MIM #603208], associated with genetic eye diseases and selected deep intronic variants were screened and analyzed as previously reported(21, 46). Variant classification followed American College of Medical Genetics and Genomics (ACMG) guidelines(47). Variants were confirmed by Sanger sequencing if variants were consistent with the phenotype, the mode of inheritance, and familial history. The datasets (variants) generated from this study were submitted to ClinVar (https://www.ncbi.nlm.nih.gov/clinvar/) (SCV001335521–SCV001335530). All patients gave written informed consent for genetic testing.

### 3. In silico analysis to predict the pathogenicity of the L144P mutation

In silico tools were used to predict the pathogenicity of L144P mutation. These were SIFT (Sorting Intolerant From Tolerant, https://sift.bii.a-star.edu.sg/www/SIFT_seq_submit2.html)(48), PolyPhen-2 (Polymorphism Phenotyping v2, http://genetics.bwh.harvard.edu/pph2/)(49), PANTHER (Protein Analysis Through Evolutionary Relationship, www.pantherdb.org) (50), SNPs & GO (https://snps.biofold.org/snps-and-go/snps-and-go.html)(51), PROVEAN (Protein Variation Effect Analyzer, http://provean.jcvi.org/index.php)(52), and PredictSNP (https://loschmidt.chemi.muni.cz/predictsnp1/)(53). SNAP-2 (screening for non-acceptable polymorphism, https://rostlab.org/services/snap2web/)(54), a neural network-based tool, was used to generate a heatmap of every possible substitution at the L144P position of Kir7.1. To predict the stability of Kir7.1 due to L144P mutation, the I-Mutant tool (https://folding.biofold.org/cgi-bin/i-mutant2.0.cgi)(55) was used. DNASTAR (Protean-3D, www.dnastar.com/)(56) and SOPMA (https://npsa-prabi.ibcp.fr/cgi-bin/npsa_automat.pl?page=/NPSA/npsa_sopma.html)(57) was used to assess differences in the biophysical properties of native and mutant protein along with the 3D graphical representation of the secondary structure.

### 4. Live-cell imaging using a heterologous expression system

HEK293 cells were cultured in a 6 well plate at 1×10^6^ cells/well (75% confluency) and 24 hours later, transfected with either pAAV-eGFP-L144P Kir7.1 or pAAV-eGFP-WT Kir7.1 plasmids using TransIT-LT1 (Mirusbio#MIR 2305). Live cell imaging was carried out after 24 hours post-transfection. The transfected cells were seeded in a 35 mm imaging dish (ibidi#81156) and stained with wheat germ agglutinin-594 (WGA-594, ThermoFisher#W11262) to label membrane, Hoechst nuclear stain (ThermoFisher#62249), endoplasmic reticulum tracker dye (ThermoFisher#E34250) were also used to assess the protein localization.

### 5. Characterization of HEK FRT WT and L144P stable cell lines

HEK Flp-In™ 293 stable cells (ThermoFisher Scientific#R75007, MA, USA) contain a single Flp Recombination Target (FRT) site at a transcriptionally active genomic locus and expresses the Zeocin^TM^ gene under SV40 early promoter. FRT site in HEK293 cells ensured the stable integration and expression of the targeted protein. These cells were maintained in complete media containing D-MEM (high glucose), 10% FBS, 1% Pen-Strep, 2mM L-glutamine, and 100 µg/ml Zeocin^TM^ for selection. According to the manufacturer’s guidelines, WT and L144P Kir7.1 HEK293 stable cell lines were created. Briefly, the FLP-In™ expression vector containing GFP tagged *KCNJ13* sequence (WT or L144P) was created by in-fusion cloning. The primers used for cloning are listed in Supplementary Table 1. The HEK293 FRT stable cells were co-transfected with pOG44 recombinase expression plasmid and FLP-In™ expression vector (Supplementary Figure 1a) containing *KCNJ13* sequence (WT or L144P). 48 hours-transfection, cells were passaged at 25% confluency for selecting stable transfectants using 400 µg/ml Hygromycin B. The Hygromycin B resistant cell clones (n=15-20) were picked, maintained in 100 µg/ml Hygromycin B and expanded for further characterization. To characterize the clones, RNA was isolated from each clone, reverse transcribed to c.DNA and subjected to Sanger sequencing using specific primers (Supplementary Table 2) to confirm the *KCNJ13* sequence (Supplementary Figure 1b). Immunocytochemistry was performed to assess protein expression and localization (Supplementary Figure 1c).

### 6. gRNA design and selection

For base editing of *KCNJ13*-L144P mutation, sgRNAs were designed using Benchling (Supplementary Figure 2a), and specific sgRNA-2 (GCTCCCAGGCCTCATGCTAG) was selected based on the highest on-target score (Supplementary Figure 2b). The chemically modified form of the sgRNA was ordered from the Synthego (CA, USA).

### 7. CRISPR-base editing of L144P mutation using C-base editors

HEK FRT stable cells expressing GFP-tagged L144P mutant proteins were subcultured for 24 hours before nucleofection at 70% confluency. The two cytosine base editor mRNAs (**1. BE4max-WTCas9**, N1me-U modification, and **2. evoCDA-SpCas9-NG**, 5moPseudoU modification) were used to edit L144P mutation along with the selected guide RNA (3:1 molar ratio). For base editing, 1×10^5^ cells were electroporated using the FS-100 program in Lonza 4D nucleofector according to the manufacturer’s guidelines. Post-electroporation, cells were seeded in a 6-well plate and maintained in 100 µg/ml Hygromycin B for further analysis.

### 8. On-target and off-target analysis using deep sequencing by Illumina platform

Five days post nucleofection, treated and untreated cells were harvested to isolate RNA (Qiagen#74134). RNA was reverse transcribed to cDNA (ThermoFisher#4368814), subsequently amplified for on-target analysis using *KCNJ13* Illumina specific primers with adapter sequences (amplicon size ~150bp). For off-target analysis, the potential off-target sites were first identified by an in-silico tool, Cas-OFFinder (http://www.rgenome.net/cas-offinder/). The parameter used were an NG/NGG/NAG PAM, with or without DNA/RNA bulge (bulge size=1) and with up to 3 mismatches to the sgRNA-2 sequence. From the treated and untreated cells, gDNA was isolated from these cells and amplified using Specific primers to generate amplicons of 150 bp. All the primer sequences are listed in Supplementary Table 3. Unique indexes (i7 and i5) were ligated to each amplicon by PCR (amplicon size 250bp), and the indexed amplicons were pooled and purified using AMPure XP beads (Beckman Coulter#A63881). The resulting library was denatured and diluted for deep sequencing on a MiSeq Illumina platform. Deep sequencing data analysis was carried out using the online tool CRISPR RGEN (http://www.rgenome.net/cas-analyzer/)(58) and CRISPResso2 (https://crispresso.pinellolab.partners.org/submission)(59).

### 9. Flow cytometry to obtain single-cell edited clones

Flow cytometry collected single cells from the pool of edited cells (Supplementary Figure 4). Single cells were grown to generate a pure clonal population of cells with different edited sequences. RNA from these cells was reverse transcribed and amplified using specific primers (Supplementary Table 2). Amplicons were subjected to Sanger sequencing using BigDye chemistry. Edited clones with either WT or synonymous changes and some nonsynonymous ones were further maintained for protein analysis and K^+^ influx.

### 10. Protein analysis by Immunocytochemistry

Kir7.1 protein expression was assessed in the mutant-L144P, WT, base-edited pooled cells, or single-cell clones by immunocytochemistry. Briefly, the cells were fixed in 4% paraformaldehyde in PBS at 4°C for 10 mins and washed twice with chilled PBS. Cells were permeabilized in 0.5% Triton X-100 in PBS (PBST) at room temperature (RT) for 5 mins and then incubated in a blocking buffer containing 2% BSA with 0.25% Triton X-100 for 2 hours at RT. As the protein is GFP tagged, GFP mouse monoclonal primary antibody (Cell Signaling#2955, 1:250) was used to detect Kir7.1 protein expression in the cells. Sodium Potassium ATPase rabbit monoclonal primary antibody (Thermo Fisher#ST0533, 1:500) was used to label the cell membranes. Primary antibody incubation was carried out at 4°C overnight. The cells were washed thrice for 5 mins/wash with chilled PBS to remove unbound primary antibodies. The cells were incubated with secondary antibodies, Alexa fluor-594 conjugated Donkey anti-Rabbit (Proteintech#SA00006.8, 1:500), and Alexa fluor-488 conjugated Donkey anti-Mouse (Proteintech#SA00006.5, 1:500) at RT for 1 hour in the dark. DAPI (Biotium#40043, 1:500) was used as a nuclear counterstain. The immunostained cells were imaged on a confocal microscope (Nikon C2 Instrument).

### 11. In silico mRNA structure prediction and half-life

RNAsnp web server (https://rth.dk/resources/rnasnp/) was used to predict the effect of single nucleotide variation on mRNA secondary structure. The cells were seeded in a 24-well plate and treated with actinomycin D (10 µg/ml) to inhibit the transcription for mRNA half-life calculation. Cells were collected at different time point (0, 0.5, 1, 1.5, 2, 3, 4 hours) for RNA isolation. RNA was reverse transcribed to cDNA, and real-time PCR was performed using SYBR green chemistry. Average Ct values of each sample at each time point were normalized to respective average Ct values of t=0 to obtain ΔCt value [ΔCt = (Average Ct of each time point - Average Ct of t=0)]. mRNA abundance for each time point was calculated by 2^(−ΔCT)^. the mRNA decay rate and half-life were determined by non-linear regression curve fitting (one phase decay) using GraphPad Prism 9.

### 12. Functional analysis of edited cells using automated patch clamp

An automated patch clamp (Q Patch II, Sophion, Denmark) measured the whole-cell current from the WT, L144P, and base edited cells. For the experiment, the cells were grown in a T75 flask for 48-72 hours and then detached gently using Detachin^TM^. The cells were centrifuged at 90 g for 1 min and resuspended in serum-free media containing 25 mM HEPES. The cells [3 M/ml] were kept on the instrument’s shaker for 20 minutes before the experiment. 48 cells were recorded in parallel using single-hole disposable Qplates through individual amplifiers. A pressure protocol was used to achieve cell positioning (−70 mbar), Giga seal (−75 mbar), and whole-cell configuration (5 pulses with −50 mbar increment between the pulses, first pulse of −250 mbar). The current was recorded in response to voltage-clamp steps from the holding potential (−10mV) to voltages between −150mV and +40mV (Δ=10mV). More than 60% of the cells completed the experiment. The cells in which the stability was compromised during the experiment were judged by the leak current and excluded from the analysis. The extracellular solution contained (in mM): 135 NaCl, 5 KCl, 10 HEPES, 10 glucose, 1.8 CaCl_2,_ and 1 MgCl_2,_ pH adjusted to 7.4 and osmolarity 305 mOsm. The intracellular solution contained (in mM) 30 KCl, 83 K-gluconate, 10 HEPES, 5.5 EGTA, 0.5 CaCl_2_, 4 Mg-ATP, and 0.1 GTP, pH adjusted to 7.2, and osmolarity 280 mOsm. For rubidium ‘ ‘Ringer’s external solution, NaCl was replaced with RbCl [140 mM] and used as an enhancer of Kir7.1 current. The data was analyzed using Sophion Analyzer v6.6.44.

### 13. tRNA sequencing

Total RNA was isolated from HEK293 FRT WT stable cells and was quantified using a NanoDrop ND-1000 instrument. The tRNAs were purified from the total RNA, demethylated, and partially hydrolyzed according to the Hydro-tRNAseq method. The tRNAs were re-phosphorylated and converted to small RNA sequencing libraries using NEBNext® Multiplex Small RNA Library Prep Set for Illumina® kit (New England Biolabs). Size selection of 140-155 bp PCR amplified fragments (corresponding to 19-35 nt tRNA fragments size range) was performed. The tRNA-seq libraries were quantified using Agilent 2100 BioAnalyzer. According to the manufacturer’s instructions, the libraries were sequenced for 50 cycles on Illumina NextSeq 500 system using NextSeq 500/550 High-Output v2 kit (75 cycles) according to the ‘ ‘manufacturer’s instructions. Sequencing quality was examined by FastQC software, and trimmed reads (pass Illumina quality filter) were aligned to cytoplasmic mature-tRNA sequences from GtRNAdb and mitochondrial tRNA sequences from mitotRNAdb using BWA software. The expression profiles of tRNAs were calculated based on uniquely mapped reads. The differentially expressed tRNAs were screened based on the count value with R package edgeR.

### 14. Statistical analysis

Each experiment was repeated three times with proper controls. Statistical analysis was performed using a two-tailed student’s t-test, and a *p*-value of 0.05 was considered statistically significant. Graphs were plotted using origin 9.1 and GraphPad Prism 9.

## Acknowledgments

We sincerely thank the affected individuals and family members for their willingness to participate in the study. We thank the Stem Cell & Regenerative Medicine Center (SCRMC) fellowship for the funding support to Meha Kabra. We acknowledge Retina Research Foundation Kathryn and Latimer Murfee Chair, McPherson ERI (KS), M.D. Matthews Research Professorship (BRP), NIH R01 EY024995 and R24 EY032434 (B.R.P.). We acknowledge the present and past members of Pattnaik lab for their valuable input.

## Supplementary

**Supplementary Table 1:**
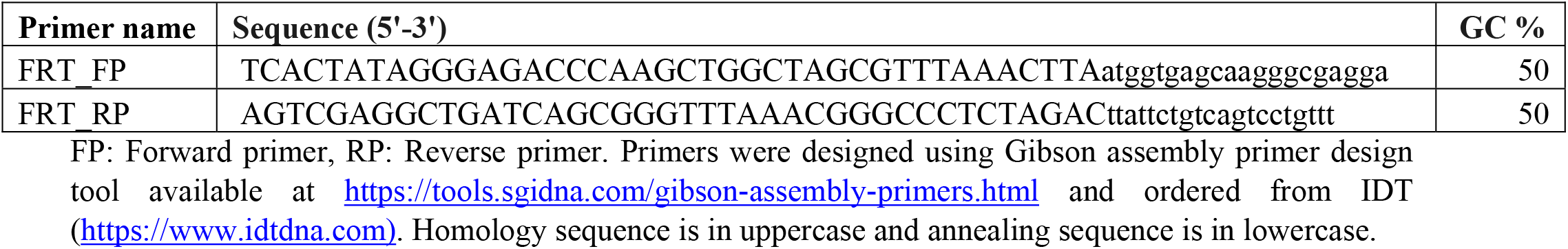
Primers for in-fusion cloning of *KCNJ13* in FLP-In™ expression vector.

**Supplementary table 2:**
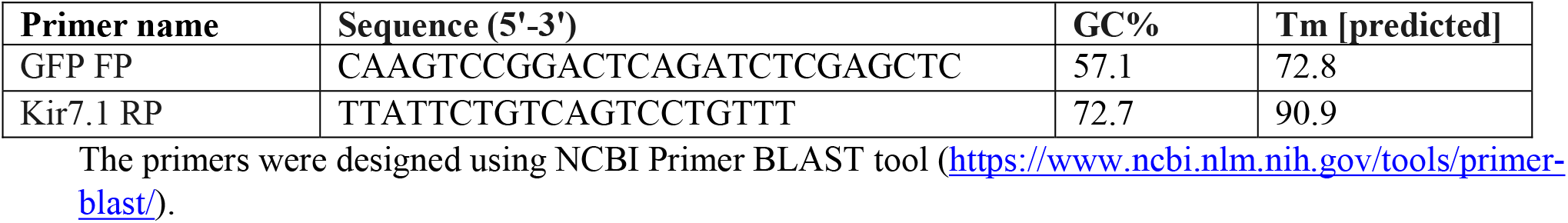
Primers for Sanger sequencing.

**Supplementary Table 3:**
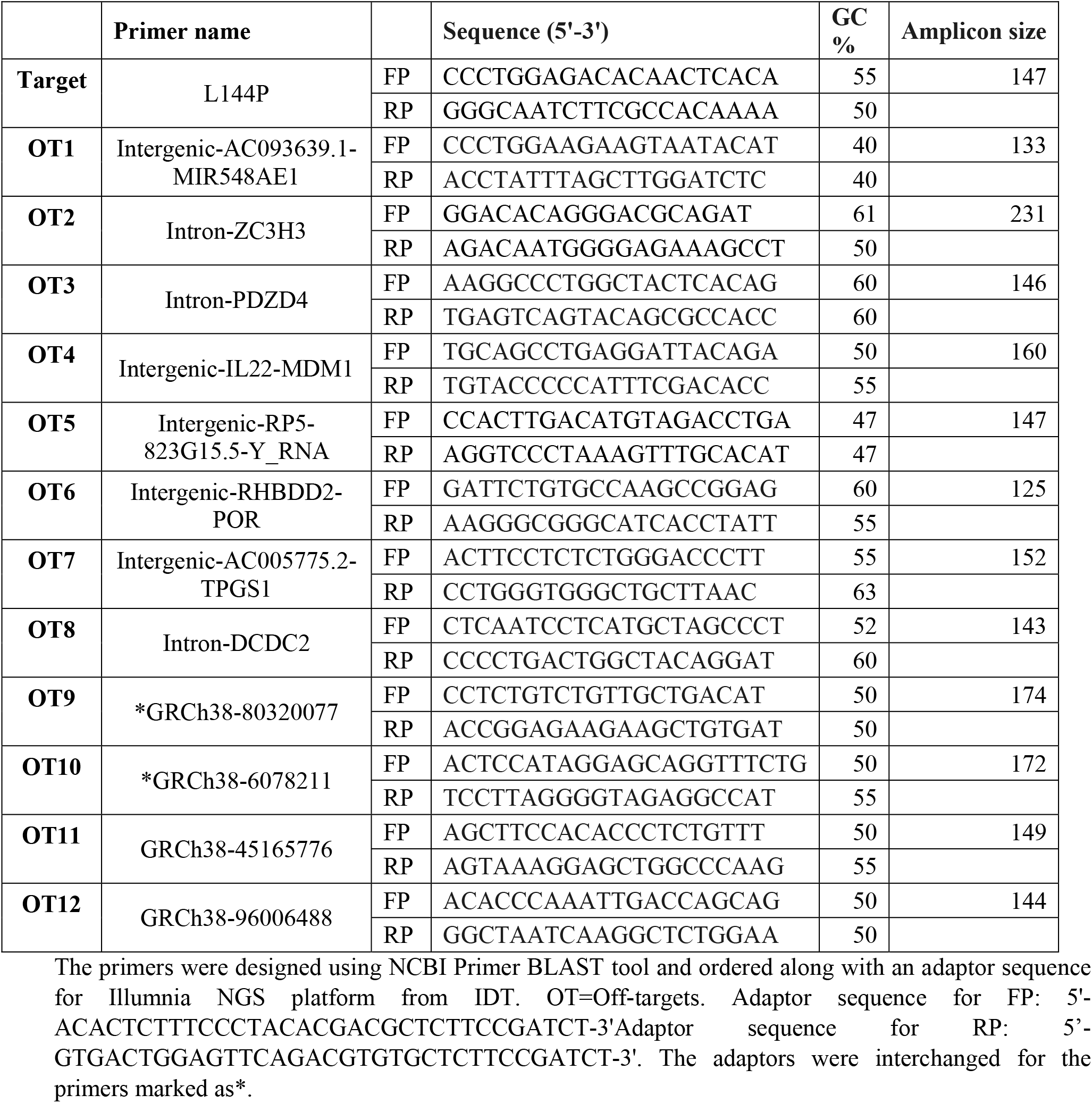
Primers to amplify the on-target and off-target sites.

**Supplementary Table 4:**
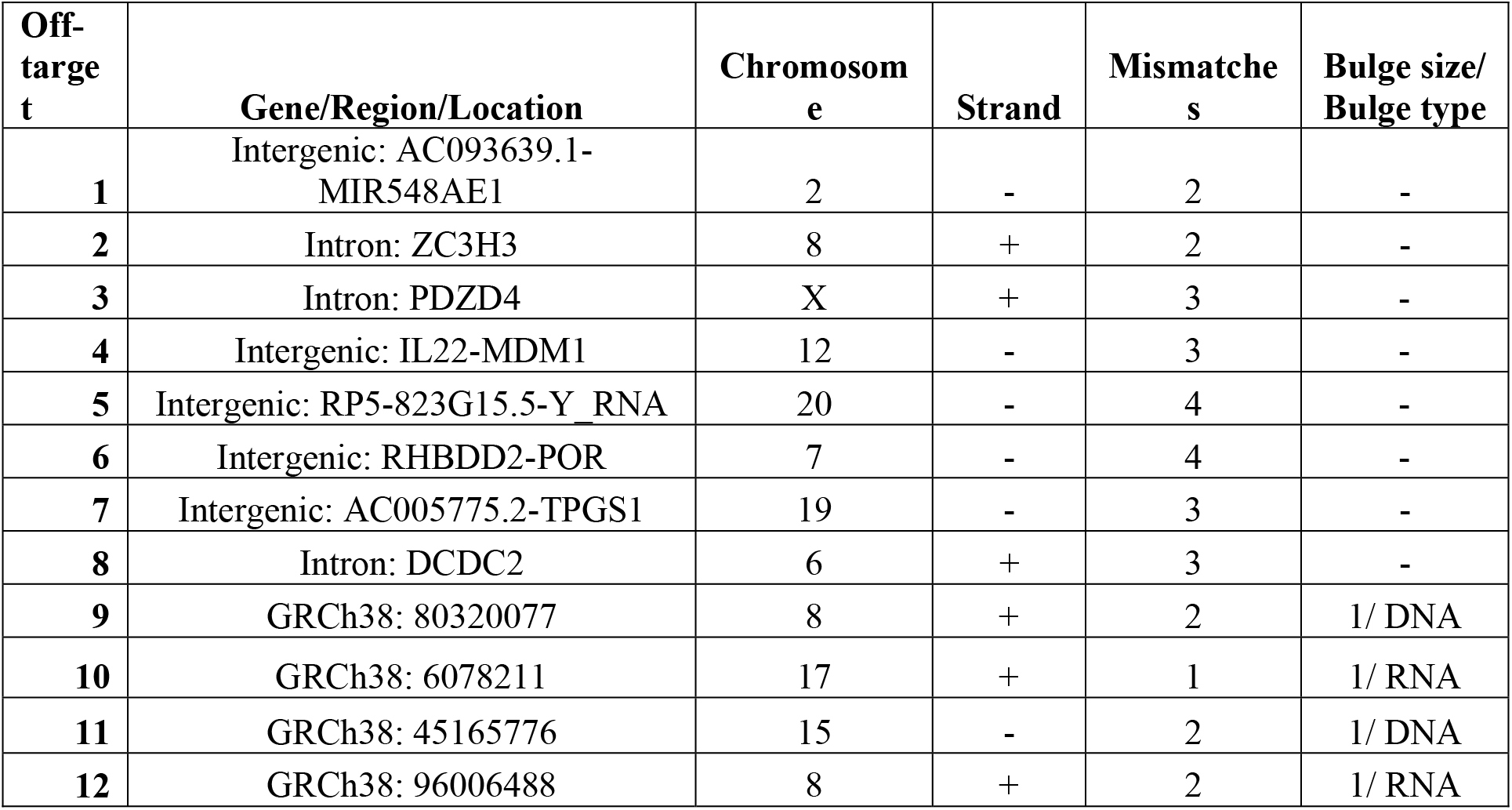
Potential off-target sites for L144P sgRNA location screened by deep sequencing.

**Supplementary Figure 1:**
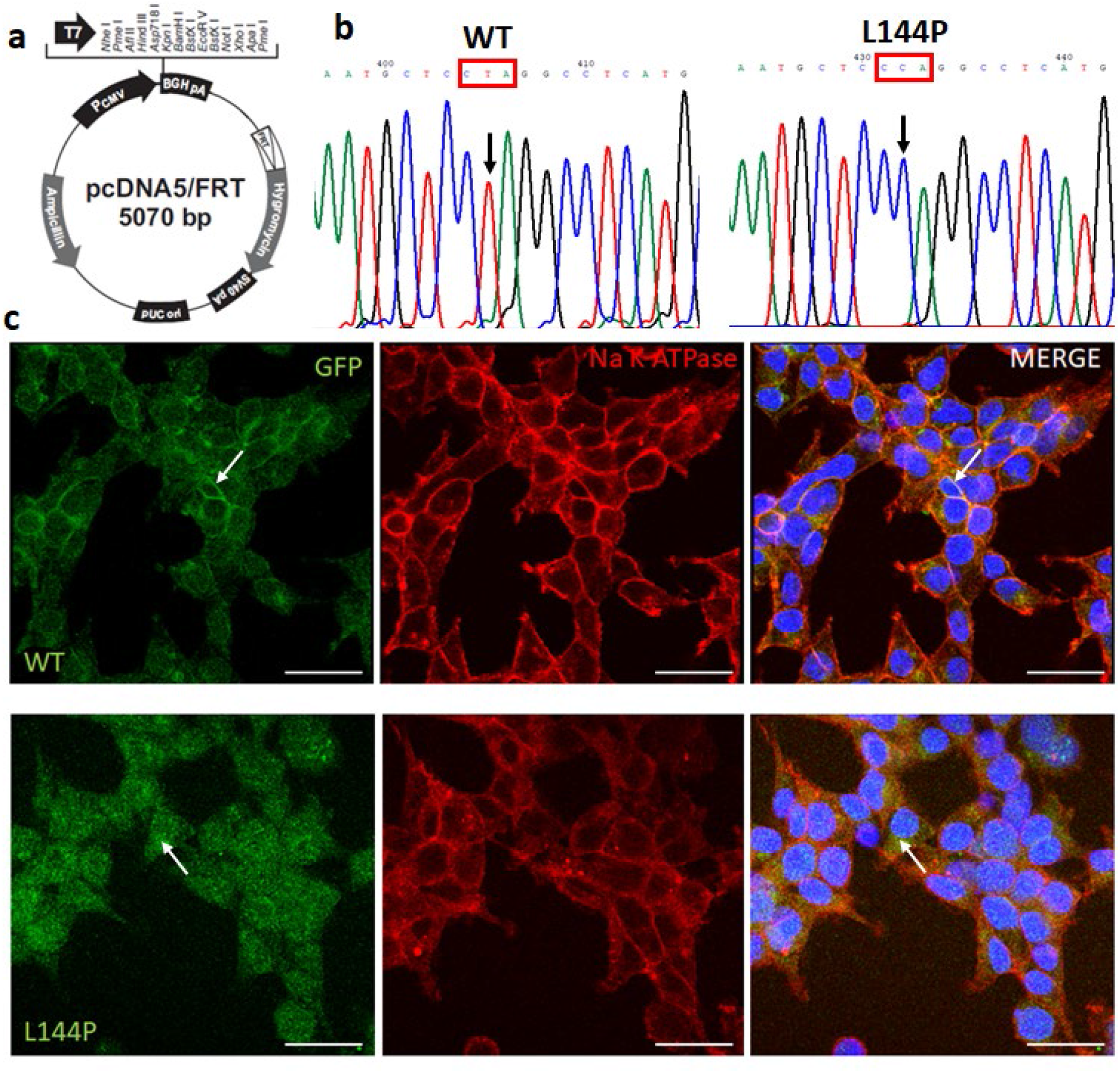
Characterization of HEK293 FRT WT and L144P stable cells. a] FLP-In™ expression vector map used for in-fusion cloning to express WT and L144P Kir7.1. b] WT and L144P mRNA sequence from respective HEK293 FRT Stable cells. c] Native and L144P Kir7.1 protein expression in stable cells assessed by immunocytochemistry.

**Supplementary Figure 2:**
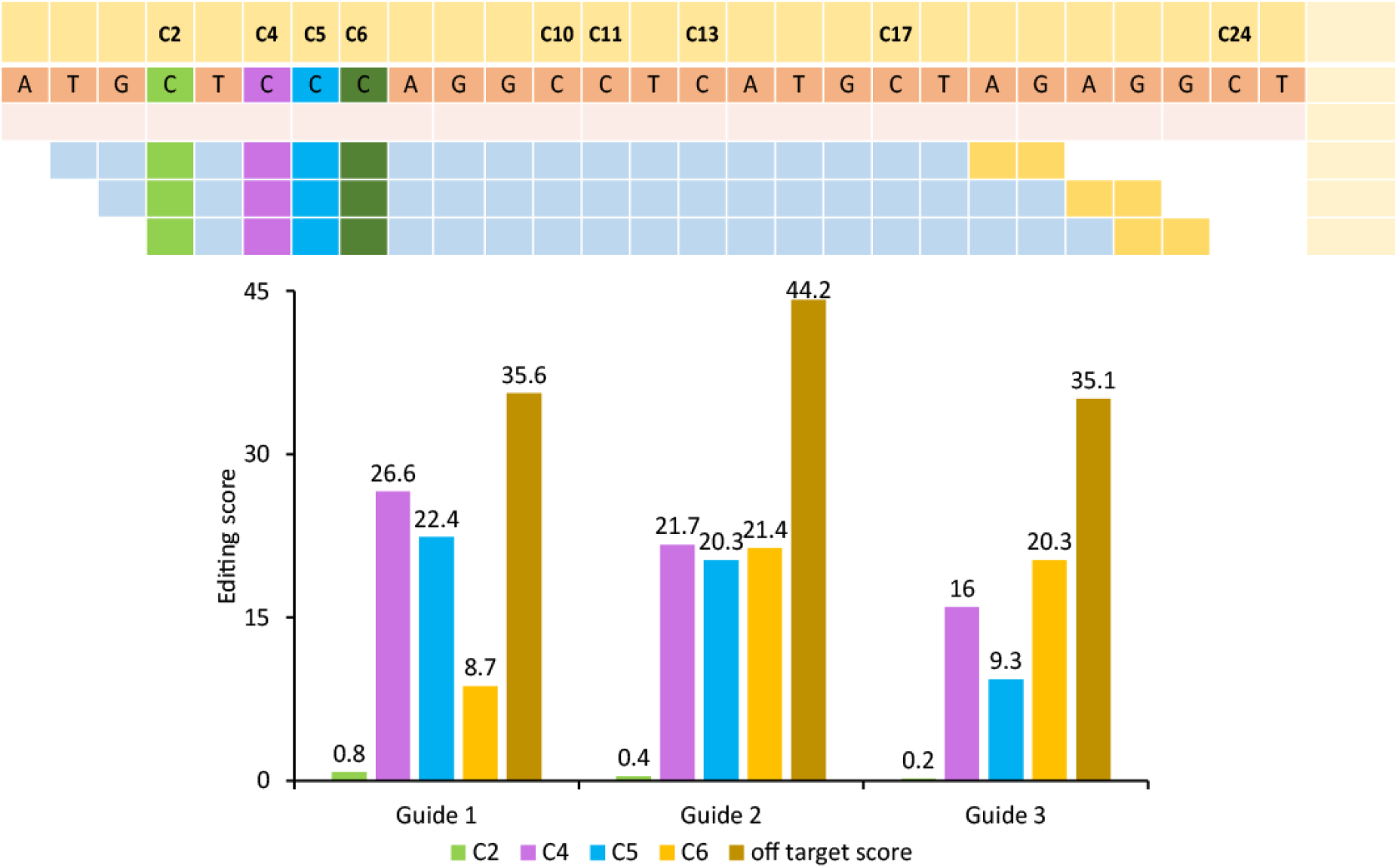
The gRNA design and selection. a] Three gRNAs with NG PAM (black-dashed rectangle) location at KCNJ13 gene sequence b] on-target scores at different neighboring Cs and off-target scores of three guides designed by Benchling software.

**Supplementary Figure 3:**
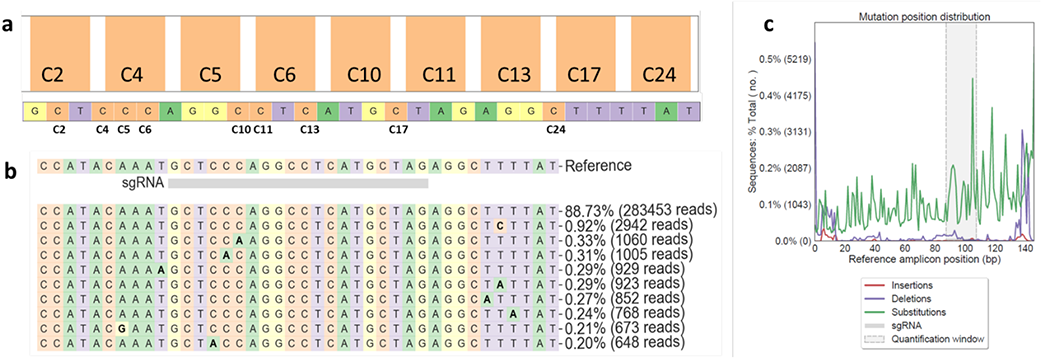
Sequencing readouts from untreated L144P stable cells used as reference. a] Nucleotide distribution around sgRNA location as observed in sequencing reads. b] Percentage of sequencing reads observed in the untreated sample. c] Percentage distribution of substitution and deletion at sgRNA location.

## Additional files-Source Data

**Figure 2 Source Data 1 (.jpg/.pptx)**

Figure 2-Source Data 1 contains the full images for Figure 2F.

**Figure 3 Source Data 1 (.pptx)**

Figure 3-Source Data 1 contains the images from different field demonstrating the localization of WT and L144P Kir7.1.

**Figure 4 Source Data 1**

Figure 4-Source Data 1 folder contains the NGS files for BE4max treated samples (n=3) in fastaq.gz format which can be analyzed using an online CRISPResso2 tool.

**Figure 4 Source Data 2**

Figure 4-Source Data 2 folder contains the NGS files for evoCDA treated samples (n=3) in fastaq.gz format which can be analyzed using an online CRISPResso2 tool.

**Figure 4 Source Data 3**

Figure 4-Source Data 3 folder contains the NGS files for untreated samples (n=3) in fastaq.gz format which can be analyzed using an online CRISPResso2 tool.

**Figure 5 Source Data 1 (.xls)**

Figure 5 Source Data1 contains the raw files generated from automated patch clamp (APC) system without excluding any data. The data from this file was filtered out based on the criteria explained in method section.

**Figure 6 Source Data 1 (.xls)**

Figure 6-Source Data 1 contains the complete list of off-targets.

**Figure 6 Source Data 2**

Figure 6-Source Data 2 folder contains the NGS files in fastaq.gz format which can be analyzed using an online CRISPR-RGEN tool.

**Figure 7 Source Data 1 (.pdf)**

Figure 7-Source Data 1 contains flow sorting images from BE4max treated cells.

**Figure 7 Source Data 2 (.pdf)**

Figure 7-Source Data 2 contains chromatograms from flow sorted single cells.

**Figure 7 Source Data 3 (.xls)**

Figure 7-Source Data 3 contains raw files generated from automated patch clamp (APC) system without excluding any data. The data from this file was filtered out based on the criteria explained in method section.

**Figure 7 Source Data 4 (.xls)**

Figure 7-Source Data 4 contains tRNA sequencing data from HEK293 FRT stable cells for Leu codon.

## References

1. Sergouniotis PI, Davidson AE, Mackay DS, Li Z, Yang X, Plagnol V, et al. Recessive mutations in KCNJ13, encoding an inwardly rectifying potassium channel subunit, cause leber congenital amaurosis. Am J Hum Genet. 2011;89(1):183–90.

2. Pattnaik BR, Shahi PK, Marino MJ, Liu X, York N, Brar S, et al. A Novel KCNJ13 Nonsense Mutation and Loss of Kir7.1 Channel Function Causes Leber Congenital Amaurosis (LCA16). Hum Mutat. 2015;36(7):720–7.

3. Khan AO, Bergmann C, Neuhaus C, Bolz HJ. A distinct vitreo-retinal dystrophy with early-onset cataract from recessive KCNJ13 mutations. Ophthalmic Genet. 2015;36(1):79–84.

4. Kabra M, Pattnaik BR. Sensing through Non-Sensing Ocular Ion Channels. Int J Mol Sci. 2020;21(18).

5. Hejtmancik JF, Jiao X, Li A, Sergeev YV, Ding X, Sharma AK, et al. Mutations in KCNJ13 cause autosomal-dominant snowflake vitreoretinal degeneration. Am J Hum Genet. 2008;82(1):174–80.

6. Galvin JA, Fishman GA, Stone EM, Koenekoop RK. Evaluation of genotype-phenotype associations in leber congenital amaurosis. Retina. 2005;25(7):919–29.

7. Fazzi E, Signorini SG, Scelsa B, Bova SM, Lanzi G. Leber’s congenital amaurosis: an update. Eur J Paediatr Neurol. 2003;7(1):13–22.

8. Yang D, Pan A, Swaminathan A, Kumar G, Hughes BA. Expression and localization of the inwardly rectifying potassium channel Kir7.1 in native bovine retinal pigment epithelium. Invest Ophthalmol Vis Sci. 2003;44(7):3178–85.

9. Shahi PK, Hermans D, Sinha D, Brar S, Moulton H, Stulo S, et al. Gene Augmentation and Readthrough Rescue Channelopathy in an iPSC-RPE Model of Congenital Blindness. Am J Hum Genet. 2019;104(2):310–8.

10. Stadtmauer EA, Fraietta JA, Davis MM, Cohen AD, Weber KL, Lancaster E, et al. CRISPR-engineered T cells in patients with refractory cancer. Science. 2020;367(6481):1001-+.

11. Lu Y, Xue JX, Deng T, Zhou XJ, Yu K, Deng L, et al. Safety and feasibility of CRISPR-edited T cells in patients with refractory non-small-cell lung cancer. Nat Med. 2020;26(5):732-40.

12. Gillmore JD, Gane E, Taubel J, Kao JT, Fontana M, Maitland ML, et al. CRISPR-Cas9 In Vivo Gene Editing for Transthyretin Amyloidosis. New Engl J Med. 2021;385(6):493–502.

13. Frangoul H, Altshuler D, Cappellini MD, Chen YS, Domm J, Eustace BK, et al. CRISPR-Cas9 Gene Editing for Sickle Cell Disease and beta-Thalassemia. New Engl J Med. 2021;384(3):252–60.

14. Kosicki M, Tomberg K, Bradley A. Repair of double-strand breaks induced by CRISPR-Cas9 leads to large deletions and complex rearrangements (vol 36, pg 765, 2018). Nature Biotechnology. 2018;36(9):899-.

15. Fu YF, Foden JA, Khayter C, Maeder ML, Reyon D, Joung JK, et al. High-frequency off-target mutagenesis induced by CRISPR-Cas nucleases in human cells. Nature Biotechnology. 2013;31(9):822-+.

16. Cradick TJ, Fine EJ, Antico CJ, Bao G. CRISPR/Cas9 systems targeting beta-globin and CCR5 genes have substantial off-target activity. Nucleic Acids Research. 2013;41(20):9584–92.

17. Aryal NK, Wasylishen AR, Lozano G. CRISPR/Cas9 can mediate high-efficiency off-target mutations in mice in vivo. Cell Death Dis. 2018;9.

18. Adikusuma F, Piltz S, Corbett MA, Turvey M, McColl SR, Helbig KJ, et al. Large deletions induced by Cas9 cleavage. Nature. 2018;560(7717):E8–E9.

19. Perez-Roustit S, Marquette V, Bocquet B, Kaplan J, Perrault I, Meunier I, et al. Leber Congenital Amaurosis with Large Retinal Pigment Clumps Caused by Compound Heterozygous Mutations in Kcnj13. Retin Cases Brief Rep. 2017;11(3):221–6.

20. Suh S, Choi EH, Leinonen H, Foik AT, Newby GA, Yeh WH, et al. Restoration of visual function in adult mice with an inherited retinal disease via adenine base editing. Nat Biomed Eng. 2021;5(2):169–78.

21. Mejecase C, Kozak I, Moosajee M. The genetic landscape of inherited eye disorders in 74 consecutive families from the United Arab Emirates. Am J Med Genet C Semin Med Genet. 2020;184(3):762–72.

22. Toms M, Dubis AM, Lim WS, Webster AR, Gorin MB, Moosajee M. Missense variants in the conserved transmembrane M2 protein domain of KCNJ13 associated with retinovascular changes in humans and zebrafish. Exp Eye Res. 2019;189:107852.

23. Tateno T, Nakamura N, Hirata Y, Hirose S. Role of C-terminus of Kir7.1 potassium channel in cell-surface expression. Cell Biol Int. 2006;30(3):270–7.

24. Nishida M, MacKinnon R. Structural basis of inward rectification: cytoplasmic pore of the G protein-gated inward rectifier GIRK1 at 1.8 A resolution. Cell. 2002;111(7):957–65.

25. Pegan S, Arrabit C, Zhou W, Kwiatkowski W, Collins A, Slesinger PA, et al. Cytoplasmic domain structures of Kir2.1 and Kir3.1 show sites for modulating gating and rectification. Nat Neurosci. 2005;8(3):279–87.

26. Yang J, Jan YN, Jan LY. Control of rectification and permeation by residues in two distinct domains in an inward rectifier K+ channel. Neuron. 1995;14(5):1047–54.

27. Pattnaik BR, Tokarz S, Asuma MP, Schroeder T, Sharma A, Mitchell JC, et al. Snowflake vitreoretinal degeneration (SVD) mutation R162W provides new insights into Kir7.1 ion channel structure and function. PLoS One. 2013;8(8):e71744.

28. Shannon M, Miller TW, Peyton B. Randolph, Mandana Arbab, Max W. Shen, Tony P. Huang, Zaneta Matuszek, Gregory A. Newby, Holly A. Rees, and David R. Liu. Continuous evolution of SpCas9 variants compatible with non-G PAMs. Nat Biotechnol. 2020;38(4):471–81.

29. Nishimasu H, Shi X, Ishiguro S, Gao L, Hirano S, Okazaki S, et al. Engineered CRISPR-Cas9 nuclease with expanded targeting space. Science. 2018;361(6408):1259–62.

30. Koblan LW, Doman JL, Wilson C, Levy JM, Tay T, Newby GA, et al. Improving cytidine and adenine base editors by expression optimization and ancestral reconstruction. Nat Biotechnol. 2018;36(9):843–6.

31. Thuronyi BW, Koblan LW, Levy JM, Yeh WH, Zheng C, Newby GA, et al. Continuous evolution of base editors with expanded target compatibility and improved activity. Nat Biotechnol. 2019;37(9):1070–9.

32. Bae S, Park J, Kim JS. Cas-OFFinder: a fast and versatile algorithm that searches for potential off-target sites of Cas9 RNA-guided endonucleases. Bioinformatics. 2014;30(10):1473–5.

33. Zuo E, Sun Y, Wei W, Yuan T, Ying W, Sun H, et al. Cytosine base editor generates substantial off-target single-nucleotide variants in mouse embryos. Science. 2019;364(6437):289–92.

34. Grunewald J, Zhou R, Garcia SP, Iyer S, Lareau CA, Aryee MJ, et al. Transcriptome-wide off-target RNA editing induced by CRISPR-guided DNA base editors. Nature. 2019;569(7756):433–7.

35. Zhou C, Sun Y, Yan R, Liu Y, Zuo E, Gu C, et al. Off-target RNA mutation induced by DNA base editing and its elimination by mutagenesis. Nature. 2019;571(7764):275–8.

36. Kim YB, Komor AC, Levy JM, Packer MS, Zhao KT, Liu DR. Increasing the genome-targeting scope and precision of base editing with engineered Cas9-cytidine deaminase fusions. Nat Biotechnol. 2017;35(4):371–6.

37. Stergachis AB, Haugen E, Shafer A, Fu W, Vernot B, Reynolds A, et al. Exonic transcription factor binding directs codon choice and affects protein evolution. Science. 2013;342(6164):1367–72.

38. Pechmann S, Frydman J. Evolutionary conservation of codon optimality reveals hidden signatures of cotranslational folding. Nat Struct Mol Biol. 2013;20(2):237–43.

39. Boel G, Letso R, Neely H, Price WN, Wong KH, Su M, et al. Codon influence on protein expression in E. coli correlates with mRNA levels. Nature. 2016;529(7586):358–63.

40. Sabarinathan R, Tafer H, Seemann SE, Hofacker IL, Stadler PF, Gorodkin J. The RNAsnp web server: predicting SNP effects on local RNA secondary structure. Nucleic Acids Res. 2013;41(Web Server issue):W475–9.

41. Komor AC, Kim YB, Packer MS, Zuris JA, Liu DR. Programmable editing of a target base in genomic DNA without double-stranded DNA cleavage. Nature. 2016;533(7603):420–4.

42. Maeder ML, Stefanidakis M, Wilson CJ, Baral R, Barrera LA, Bounoutas GS, et al. Development of a gene-editing approach to restore vision loss in Leber congenital amaurosis type 10. Nat Med. 2019;25(2):229–33.

43. Mauger DM, Cabral BJ, Presnyak V, Su SV, Reid DW, Goodman B, et al. mRNA structure regulates protein expression through changes in functional half-life. Proc Natl Acad Sci U S A. 2019;116(48):24075–83.

44. Gehrke JM, Cervantes O, Clement MK, Wu Y, Zeng J, Bauer DE, et al. An APOBEC3A-Cas9 base editor with minimized bystander and off-target activities. Nat Biotechnol. 2018;36(10):977–82.

45. Chen PJ, Hussmann JA, Yan J, Knipping F, Ravisankar P, Chen PF, et al. Enhanced prime editing systems by manipulating cellular determinants of editing outcomes. Cell. 2021;184(22):5635–52 e29.

46. Patel A, Hayward JD, Tailor V, Nyanhete R, Ahlfors H, Gabriel C, et al. The Oculome Panel Test: Next-Generation Sequencing to Diagnose a Diverse Range of Genetic Developmental Eye Disorders. Ophthalmology. 2019;126(6):888–907.

47. Richards S, Aziz N, Bale S, Bick D, Das S, Gastier-Foster J, et al. Standards and guidelines for the interpretation of sequence variants: a joint consensus recommendation of the American College of Medical Genetics and Genomics and the Association for Molecular Pathology. Genet Med. 2015;17(5):405–24.

48. Ng PC, Henikoff S. SIFT: predicting amino acid changes that affect protein function. Nucleic Acids Research. 2003;31(13):3812–4.

49. Adzhubei IA, Schmidt S, Peshkin L, Ramensky VE, Gerasimova A, Bork P, et al. A method and server for predicting damaging missense mutations. Nat Methods. 2010;7(4):248–9.

50. Tang H, Thomas PD. PANTHER-PSEP: predicting disease-causing genetic variants using position-specific evolutionary preservation. Bioinformatics. 2016;32(14):2230–2.

51. Capriotti E, Calabrese R, Fariselli P, Martelli PL, Altman RB, Casadio R. WS-SNPs&GO: a web server for predicting the deleterious effect of human protein variants using functional annotation. BMC Genomics. 2013;14 Suppl 3:S6.

52. Choi Y, Chan AP. PROVEAN web server: a tool to predict the functional effect of amino acid substitutions and indels. Bioinformatics. 2015;31(16):2745–7.

53. Bendl J, Stourac J, Salanda O, Pavelka A, Wieben ED, Zendulka J, et al. PredictSNP: robust and accurate consensus classifier for prediction of disease-related mutations. PLoS Comput Biol. 2014;10(1):e1003440.

54. Yachdav G, Hecht M, Pasmanik-Chor M, Yeheskel A, Rost B. HeatMapViewer: interactive display of 2D data in biology. F1000Res. 2014;3:48.

55. Capriotti E, Fariselli P, Casadio R. I-Mutant2.0: predicting stability changes upon mutation from the protein sequence or structure. Nucleic Acids Res. 2005;33(Web Server issue):W306–10.

56. (TM). P. 3D Version 17.3. DNASTAR. Madison, WI.

57. Geourjon C, Deleage G. SOPMA: significant improvements in protein secondary structure prediction by consensus prediction from multiple alignments. Comput Appl Biosci. 1995;11(6):681–4.

58. Bae S. PJ, Kim J.-S. Microhomology-based choice of Cas9 nuclease target sites. Nat Methods. 2014;11:705–6.

59. Clement K RH, Canver MC, Gehrke JM, Farouni R, Hsu JY, Cole MA, Liu DR, Joung JK, Bauer DE, Pinello L. CRISPResso2 provides accurate and rapid genome editing sequence analysis. Nat Biotechnol. 2019:37(3):224–6.

